# Spatial Transcriptomic Sequencing of a DIPG-infiltrated Brainstem reveals Key Invasion Markers and Novel Ligand-Receptor Pairs contributing to Tumour to TME Crosstalk

**DOI:** 10.1101/2024.05.07.593050

**Authors:** Anja Kordowski, Onkar Mulay, Xiao Tan, Tuan Vo, Ulrich Baumgartner, Mellissa K. Maybury, Timothy E. G. Hassall, Brandon J. Wainwright, Lachlan Harris, Quan Nguyen, Bryan W. Day

## Abstract

Emerging spatially-resolved sequencing technologies offer unprecedented possibilities to study cellular functionality and organisation, transforming our understanding of health and disease. The necessity to understand healthy and diseased tissues in its entirety becomes even more evident for the human brain, the most complex organ in the body. The brain’s cellular architecture and corresponding functions are tightly regulated. However, when intercellular communications are altered by pathologies, such as brain cancer, these microenvironmental interactions are disrupted.

DIPG is a brainstem high-grade glioma arising in young children and is universally fatal. Major disease obstacles include intratumoural genetic and cellular heterogeneity as well as a highly invasive phenotype. Recent breakthrough studies have highlighted the vital oncogenic capacity of brain cancer cells to functionally interact with the central nervous system (CNS). This CNS-crosstalk crucially contributes to tumour cell invasion and disease progression. Ongoing worldwide efforts seek to better understand these cancer-promoting CNS interactions to develop more effective DIPG anti-cancer therapies.

In this study, we performed spatial transcriptomic analysis of a complete tumour-infiltrated brainstem from a single DIPG patient. Gene signatures from ten sequential tumour regions were analysed to assess disease progression and to study DIPG cell interactions with the tumour microenvironment (TME). We leveraged this unique DIPG dataset to evaluate genes significantly correlated with invasive tumour distal regions versus the proximal tumour initiation site. Furthermore, we assessed novel ligand-receptor pairs that actively promote DIPG tumour progression via crosstalk with endothelial, neuronal and immune cell communities, which can be utilised to support future research efforts in this area of high unmet need.

## INTRODUCTION

Diffuse Intrinsic Pontine Glioma (DIPG), the most common subtype of H3 K27-altered diffuse midline Glioma (DMG), is an aggressive paediatric high-grade glioma with no curative treatment options. These tumours affect children at an approximate median age of 6 to 7 years, with the majority succumbing to disease within one year [1, 2]. The development of promising targeted therapies for DIPG has long been hindered due to the lack of tumour tissue for laboratory research. Major efforts initiated by Monje and colleagues, established the first DIPG cell line and PDX animal model from patient autopsy tissue [3]. This success led to broad access to DIPG biopsy and autopsy specimens, and formed the basis of global research efforts that continue to grow today [3]. Major findings in the field include the identification of loss of histone H3 trimethylation at lysine 27 (H3K27me^3^), which is driven by somatic point mutations in genes encoding for histone H3.1 (*HIST1H3B*/*C*) and H3.3 (*H3F3A*) or overexpression of EZH inhibitory protein (*EZHIP*) [4–6]. Additional subtype-specific and co-occurring alterations frequently observed in these tumours include mutations in tumour-suppressor genes such as *TP53* or *PPM1D* or cancer-signalling genes like *PIK3CA* and *ACVR1* as well as *PDGFRA* or *EGFR* amplification/mutation [7–10]. These findings subsequently led to crucial transformations as to how DIPG was treated and classified [11]. Beyond the mutational landscape, Filbin and colleagues shed light onto tumour cell-intrinsic properties and developmental programs of H3 K27-altered gliomas. Comprehensive sequencing analyses at the single cell level revealed that DMG tumours are largely comprised of oligodendrocyte precursor (OPC-like) cells that give rise to more differentiated astrocyte (AC-like)/mesenchymal (MES-like) and oligodendrocyte (OC-like) cell populations. Extended work by Liu et al. have shown that OPC-like populations can be further divided into more immature (OPC-like 2 and 3) and more committed OPC-like 1 states [12, 13]. Additionally, the group examined the architecture of these tumours at the single-cell spatial level, resolving cellular relationships between populations and suggesting the presence of genetically and functionally distinct tumour niches.

Recent breakthrough studies uncovered that glioma cells, including DIPG, take advantage of intercellular communication and actively integrate into neuronal networks. Pioneering work in the emerging field of Cancer Neuroscience sought to elucidate how crosstalk with neurons and neuronal activity regulate tumour growth and invasion. Such interactions crucially promote glioma pathogenesis via non-synaptic mitogenic paracrine mechanisms as well as through neuron-to-glioma synapse electrochemical signalling [14–18]. In the context of adult glioblastoma (GBM), Winkler and colleagues unravelled the complexity of these networks further, demonstrating that intercellular communication is cell type-specific and that different glioma subpopulation functionally connect with each other in different ways [19]. Collectively, these discoveries emphasise the intricacy of spatiotemporal heterogeneity present in these tumours, highlighting the unfavourable impact of the interconnectome that exists between tumour cells and the microenvironment.

In collaboration with the Queensland Children’s Tumour Bank (QCTB) in Brisbane, we collected fresh tumour tissue from a single DIPG patient at autopsy, spanning the entire tumour-infiltrated brainstem from midbrain down to the spinal cord. Whilst numerous genetic studies on DIPG tumours from spatially distinct locations have been reported, analysis on wholly intact tissue remains sparse. To address this unmet need, we performed 10x Visium spatial transcriptomic sequencing on 10 sequential DIPG brainstem regions. Our analysis has revealed additional meaningful insights into the molecular mechanisms and novel ligand-receptor cell interactions that actively support DIPG progression and crosstalk with the TME.

### CASE REPORT AND SAMPLE OVERVIEW

A boy, aged 7 years and 7 month, presented as part of a general clinical investigation for a unilateral coloboma, with an incidental finding of a T2 and FLAIR hyperintense lesion (AP: 25mm, TP: 22mm, CC: 35mm) involving the inferior pons and medulla dorsally (**Figure 1A.I**). The patient displayed no specific signs of neurological deficit and remained well for a period of 6 years and 10 months with no imaging changes or indication of disease progression. 7.5 years after the initial scan, he had a pre-syncopal episode and reported new symptoms including headaches, increased slurring of speech and ataxia. MRI showed an infiltrative neoplastic lesion, demonstrating moderate interval progression of the pons and medulla with new involvement of the cerebellar peduncles (**Figure 1A.II**). Histopathology and molecular profiling from stereotactic biopsy sampling diagnosed a diffuse midline glioma, H3 K27M mutant (*H3F3A* c83.A>T), MGMT unmethylated, with *TP53*, *ATRX* and *PPMD1D* somatic point mutations and copy number variation loss of chromosome 5q, 15q and 18 (**Table Figure 1**). The patient received standard of care local field radiotherapy (RT) of 50 Gy in 30 fractions using a VMAT technique, resulting in no clinical improvement. MR imaging 5 month post RT demonstrated concerning disease progression (35 x 43 x 65 mm), involving all cerebellar peduncles as well as the anterior aspect of the right cerebellar hemisphere extending further superiorly. There was additional interval increase in size of the lateral and third ventricles and cerebellar tonsillar descent below the foramen magnum (**Figure 1A.III**). The patient remained on dexamethasone and received palliative care. He passed away 9 weeks after the last scan and the family generously donated his sample through the QCTB.

**Figure 1:**
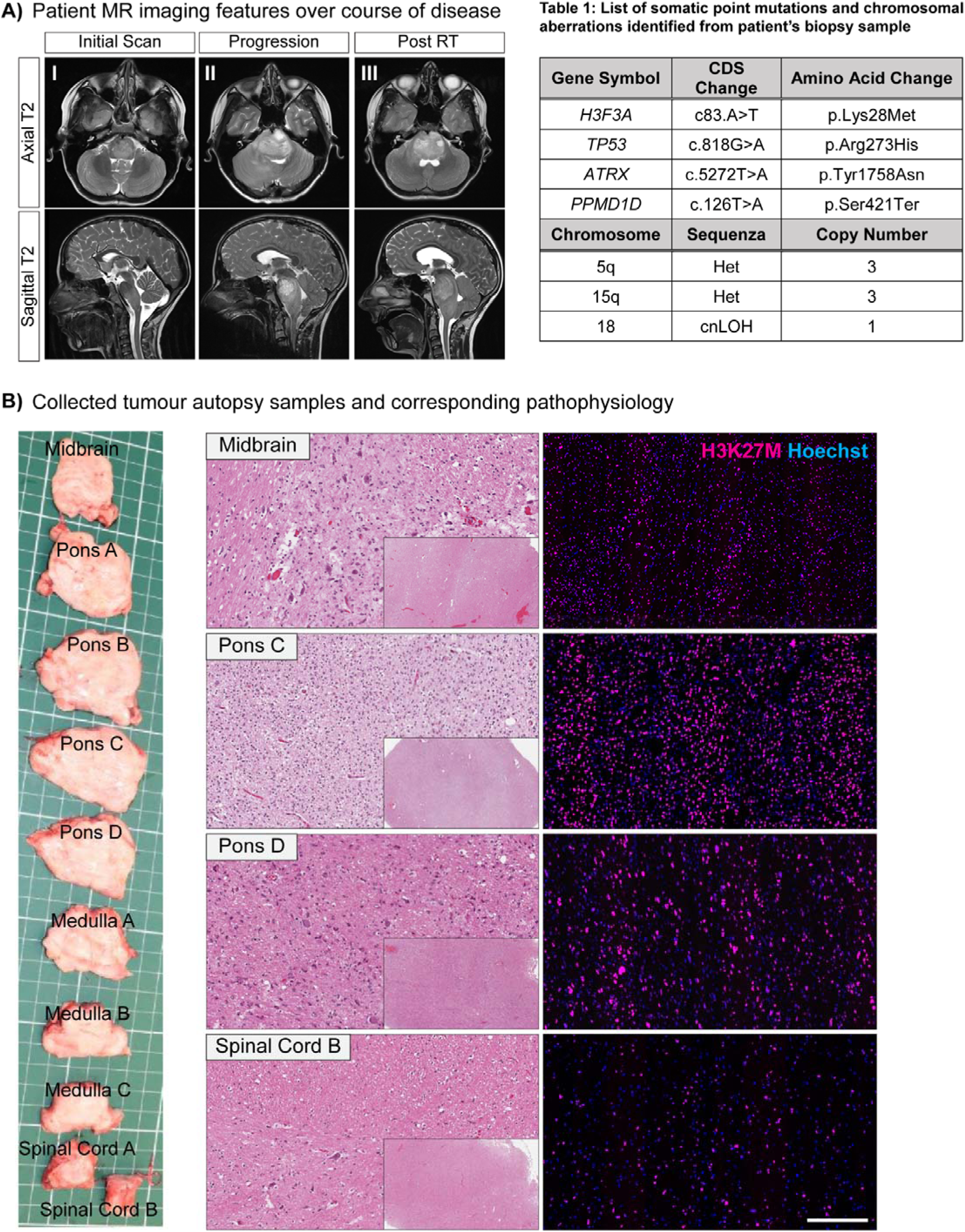
Patient imaging and tumour characteristics. **(A)** MR images showing DIPG tumour progression over the course of initial presentation to disease end stage. Table summarising point mutations and chromosomal aberrations identified through tumour biopsy sampling and sequencing as part of the MNP2.0 (Germany) and PRISM (Sydney) trial. **(B)** 10 patient brainstem regions were collected and different degrees of tumour burden confirmed by H&E and H3K27M staining. Pontine regions showing high cellularity areas, whereas clear patterns of tumour infiltration are observed in the midbrain area.

Research autopsy was performed within hours of the patient’s death and tissue annotated and banked. The diseased brainstem was segmented into ten cross-sections, including midbrain, pons, medulla and spinal cord. Formalin-fixed and paraffin-embedded (FFPE) tissue blocks were prepared for each of the ten samples and assessed for tumour burden. H&E staining demonstrated varying degrees of neoplastic cells, with high cellularity areas in pontine regions and obvious patterns of tumour infiltration in the midbrain. The extent of DIPG tumour burden was confirmed via H3K27M immunohistochemistry (IHC) on matching FFPE tissue blocks (**Figure 1B and Supplementary Figure S1A**). RNA quality (DV200 values) and the degree of H3K27M-positive cells was determined for all FFPE tissue blocks, confirming suitability of the samples for sequencing studies (**Supplementary Figure S1B**).

## RESULTS

### Spatially informed cell type clustering highlights heterogeneous cell community distribution

Quality control and filtering was performed on each region individually (**Supplementary Figure S2A**). To further compare the 10 different brainstem regions, the sequencing data from all samples was inferred jointly via anchor-based data integration. This step was employed to minimise batch effects and allow for the identification of conserved and shared features across all samples [20]. Visualisation of all the regions in Uniform Manifold Approximation and Projection (UMAP) confirmed successful data integration (**Supplementary Figure S2B**). Unsupervised cell type clustering was performed on the integrated dataset, resulting in ten unique populations, herein referred to as cell communities. The ten communities represent individual spots, dominated by the following brain cell types: stem- and progenitor-like, glial, neuronal, myeloid and endothelial cell populations (**Figure 2B, C**). Proportional quantification of these further showed that communities with an OPC/NPC (C0) and stem-like/glial (C1) phenotype were the dominating cell populations in our dataset (**Figure 2B**). Spatial profiling and visualisation demonstrated a diffuse distribution of different cell communities in all tissue regions (**Figure 2C and Supplementary Figure S2C**). Certain communities, including those with stem- and progenitor features, were in turn sparsely expressed, hinting at distinct subpopulations with specific functions. Liu et al. have previously analysed the spatial architecture of four H3 K27M DMG patient samples, performing IF-based co-detection by indexing (CODEX) [13]. Similar to the results from this group, we found that the ten cell communities show varying clustering tendencies in a cell type-dependent manner. The OPC/NPC, stem-like/glial as well as neural populations predominantly cluster together with themselves, whereas myeloid cells or mature astrocytes/oligodendrocytes appeared to be more dispersed in the majority of the ten brainstem regions. Interestingly, myeloid and neural communities largely surround the larger OPC/NPC and stem-like/glial populations indicating actively infiltrating immune cells and suggesting areas of tumour-CNS interactions. Overall, this spatially-resolved delineation of the tumour-infiltrated brainstem highlights the complex cellular heterogeneity present in this disease and suggests distinct cellular interactions.

**Figure 2:**
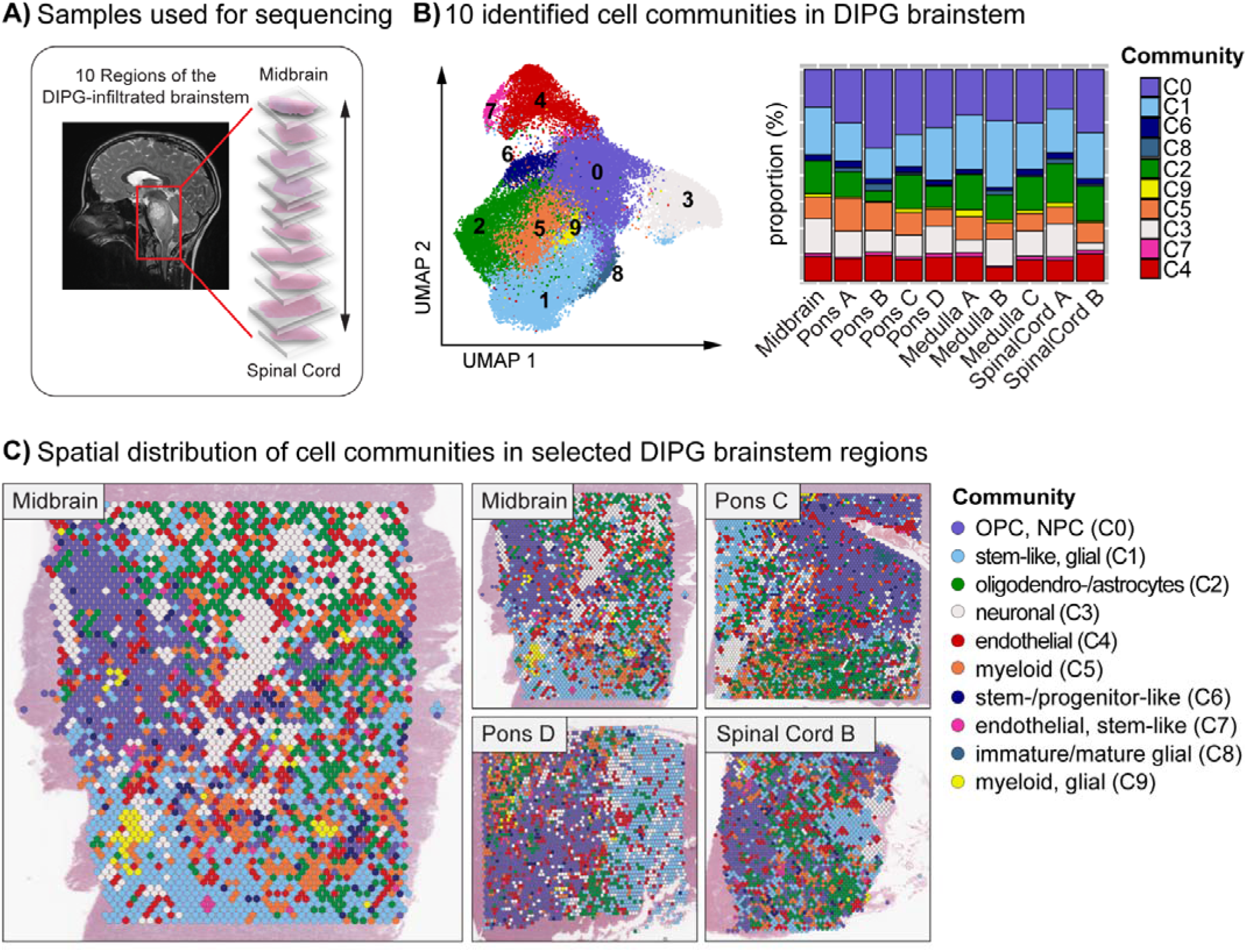
Identification cell communities in the DIPG-infiltrated brainstem. **(A)** Schematic representation of sequenced DIPG-infiltrated brainstem regions. **(B)** Unsupervised cell type clustering leading to the identification of 10 cell communities. Corresponding proportions highlighting community C0 and C1 as the dominating populations in all brainstem regions. **(C)** Spatial visualisation of the 10 cell communities in the brainstem regions showing heterogeneous distribution of the respective populations, including patterns of immune cell infiltration and highly compartmentalised OPC or stem-like populations.

### Identification of tumour cell communities underlines co-existing DIPG subpopulations

Comprehensive sequencing studies from Suvà, Filbin and colleagues have identified the cellular origin and hierarchies underlying DMG tumourigenesis. Findings show that these tumours predominantly resemble cells of OPC and stem-like origin that give rise to more differentiated oligo- or astroglial populations [12, 13]. Hence, it was reasonable to hypothesise that cell communities with OPC, NPC and stem-like phenotypes, in our dataset, could represent regions of higher tumour content. To assess this this computationally, we performed InferCNV, an established method to identify evidence for large-scale copy number variations (CNV) in single cell and spatial tumour sequencing data [21]. To further support our data, we incorporated three normal, human dorsolateral prefrontal cortex 10x Genomics Visium spatial transcriptomics datasets, derived from the web application spatialLIBD [22]. Amongst others, we found CNVs that were identified via biopsy sampling, including loss of chromosome (Chr) 5 and 15, as well as those characteristic for DMG tumours, such as gain of Chr1q [23]. Chr1 gain and Chr15 loss were present consistently across all ten regions, but were absent in non-malignant normal brain tissue (**Figure 3A**). To identify which cell populations contain a high degree of the identified chromosomal aberrations, we determined the expression of the CNVs in the ten cell communities. Results confirmed that predominantly populations with an OPC and stem- or progenitor-like phenotype harbour aberrations in the defined chromosomes. These included the strongly represented OPC/NPC (C0) and stem-like/myeloid (C1) cell communities, as well as the modestly populations of C6 and C8 (**Figure 3B**). The identification of four ‘tumour cell communities’ raised the question whether these are characterised by genetic differences and represent distinct transcriptional profiles. Previous work form Vinci et al. discovered that DIPG is comprised of numerous distinct subclones that co-exist and demonstrated that such cell populations are genetically and functionally diverse. Following studies provided further evidence that these subclonal populations likely interact co-operatively to maintain tumourigenesis and promote a DIPG invasive phenotype in a spatiotemporal context [24–27]. Next, we therefore sought to delineate patterns of heterogeneous DIPG tumour populations and assessed expression levels of selected marker genes associated with DIPG cell signatures. Results indicated that the four malignant populations reflect distinct tumour subclones (**Figure 3C**). For instance, C0 contains well-known DIPG-associated genes, such as *NTRK2*, *EGFR*, or *OLIG2*. This community further displayed gene signatures that are correlated to the more committed OPC-like1 cellular state, including *EPN2* or *POU3F3*. Community C1, in turn, showed elevated stem cell and regulatory genes like *JUNB* and *FOS,* which assign to the more immature OPC-like3 state [13]. Community C6, which appears more widely dispersed across the respective tissue sections, is predominantly characterised by developmental and cell cycle genes including *MKI67* or *CDC20* among others. C8, which is represented to minor extent, strongly express a combination of genes between C0 and C1, but additionally harbours DNA damage response genes, such as *DDIT4*. To explore this further and to identify potential relationships between these tumour subpopulation, we performed spatial trajectory analysis. Supporting our hypothesis, results suggested a trajectory tree from immature to more committed DIPG cells, proposing a transition from C6 to C1 to C8 to C0 (**Figure 3D and Supplementary Figure S3A**). As described above, spatially-resolved visualisation of the four tumour cell communities showed that the communities C8 and C0 largely occur compartmentalised, whilst C6 appears scattered across the tissue sections. C1 primarily clusters with itself, but is more widely distributed compared to C0 and C8 (**Figure 3D and Supplementary Figure S3B**). Hence, this finding strongly supports previous data, illustrating the presence of functionally distinct tumour niches in a spatiotemporal context and suggests transitioning from immature to more committed DIPG tumour cell populations.

**Figure 3:**
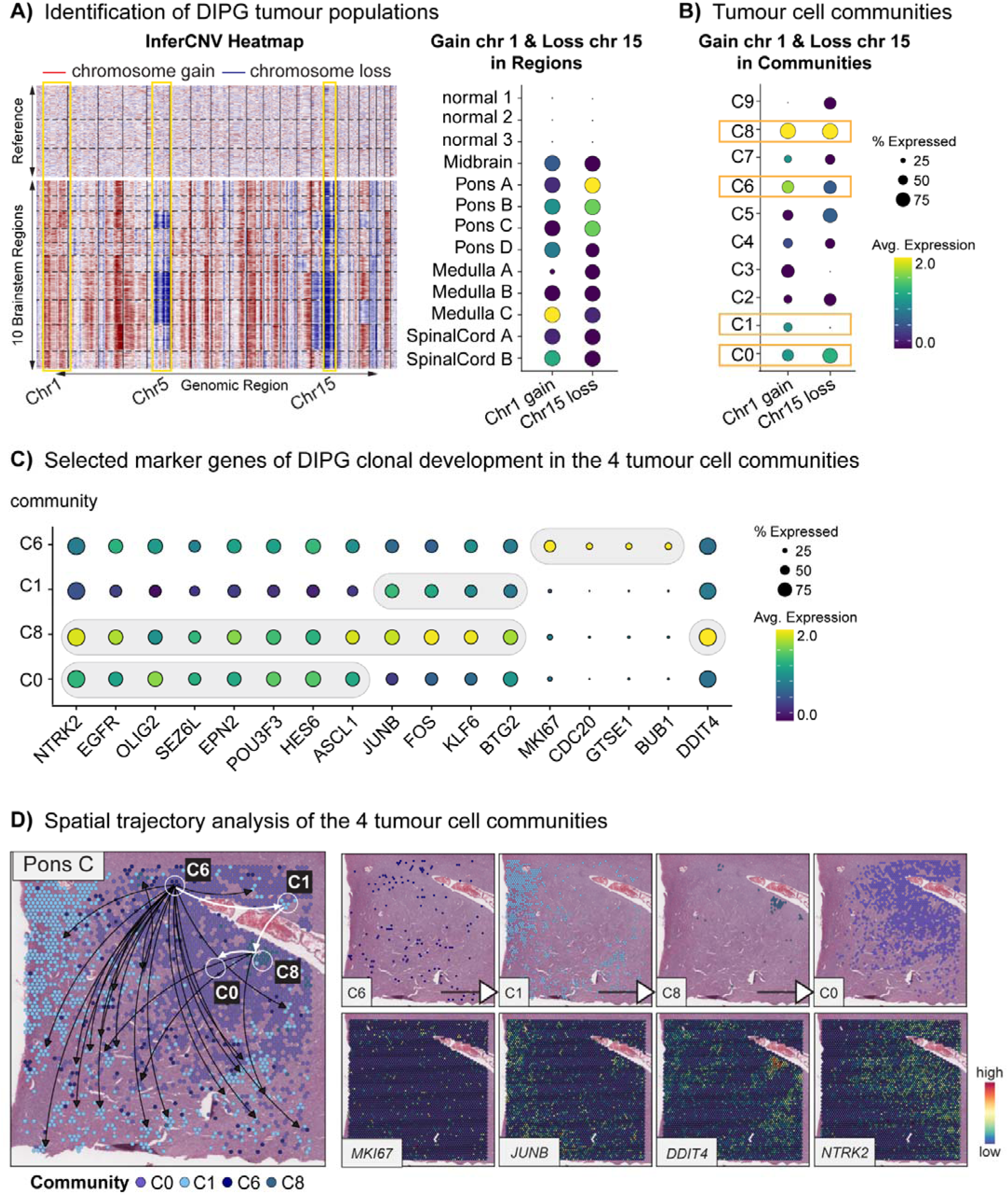
Identification and characterisation of tumour cell communities. **(A)** InferCNV analysis of the 10 DIPG brainstem regions and 3 normal references showing chromosomal aberrations of chr1, chr5 and chr15 in the tumour-infiltrated regions. Expression profiling of chr1 gain and chr15 loss confirming these aberrations are present in the sequenced DIPG samples but not in the normal reference brain. **(B)** CNV profiling grouped by cell communities reveals that predominantly C0, C1, C6 and C8 harbour gain of chr1 and loss of chr15. **(C)** The levels of selected marker genes according to Liu et al. were assessed across the identified tumour cell communities, showing that C0 expresses known DIPG markers like *NTRK2* or *EGFR*, whereas C1 demonstrates genes associated with immature OPC-like populations such as *JUNB*. C6 expresses proliferation markers like *MKI67* and C8 appears to be a mix of C0 and C1, with additional DNA damage markers. **(D)** Spatial trajectory interference proposing clonal development from C6 to C1 to C8 to C0, supporting a trajectory path from more immature to more committed DIPG tumour cells.

### Genetic profiling of tumour-distal regions highlights neuronal gene signatures

One hallmark of DIPG is the diffuse and highly infiltrative nature of these tumours. Our dataset offered the opportunity to assess the genetic profile of areas that are more strongly correlated with the tumour invasive edge. Hence, we next compared tumour distal areas that demonstrate clear patterns of tumour infiltration to the tumour-initiating proximal regions. For this, we determined the expression levels of all 18,025 genes in all brainstem regions and further calculated the Pearson correlation coefficient (r), quantitating gene expression changes from region Medulla A towards the Midbrain or Pons D towards Spinal Cord B. Our analysis highlighted 94 genes that were significantly (p>0.049) associated with distal areas and demonstrated high correlation values, ranging between 0.81 (*GAS7*) and 0.98 (*POU4F1*). **Figure 4A** visualises the magnitude of expression of these genes in order of r ranking. Interestingly, many of these genes reflect neuronal gene signatures. The importance of neuronal activity to glioma invasion, especially DIPG, has been well established in recent years [14–18, 28–31]. *POU4F1*, the most strongly correlated gene with tumour-distal regions, belongs to the POU (Pit-Oct-Unc) class of neural transcription factors and critically determines cell fate of somatosensory neurons and maintenance of mature neurons [32, 33]. *PTGER3*, the second highest correlated gene, encodes for prostaglandin receptor E3. Prostaglandin receptors have been shown to be expressed on excitatory neurons and to interact with prostaglandin E2 secreting microglia, thereby modulating glycineric synapses [34]. Similarly, two members of the complement system, *C3* and *C1QA*, are amongst the 94 genes. Microglia and neurons interact through C3 and C1Q during nervous system development, which is known to be a potent “eat me’ signal and established to regulate synapse pruning [35, 36]. Compared to *POU4F1* or *PTGER3*, *C3* and *C1QA* are comparatively highly expressed in our dataset, underscoring important involvement of the immune cell population in brain pathophysiology. We performed KEGG and Gene Ontology (GO) analysis of the 94 genes, confirming that the transcriptional programs were indeed highly associated with neuroactive ligand-receptor interactions and immune signatures/the complement cascade. These appeared to be mainly regulated via cAMP (cyclic adenylyl cyclase monophosphate) and G protein-coupled receptors (GPCR) signalling pathways (**Figure 4B**). Of note, (nerve growth factor inducible) *VGF* was both highly expressed and highly correlated with distal regions. This peptide has been previously demonstrated to be a critical regulator for BDNF-TrkB signalling [37, 38]. The importance of this pathway contributing to neuron-to-glioma synapse formation and plasticity was highlighted only recently by Taylor and colleagues [39]. We also observed *TIMP1* (Tissue inhibitor matrix metalloproteinase 1) highly expressed and positively correlated with distal regions. This protein is a prognostic marker in adult glioblastoma (GBM) and has been linked to chemoresistance, likely via TIMP1-CD63-mediated stem cell maintenance [40]. Amongst all 94 genes, *DTNA* (dystrobrevin alpha/ αDG) was found highest expressed and elevated in almost all brainstem regions. This member of the dystrophin family is predominantly found within perivascular astrocyte endfeet and neurons [41]. Notably, the alpha isoform of dystrobrevin in particular was attributed a crucial role for neuronal migration and synaptic transmission during development [42]. Aside from these examples, two other candidates were significantly correlated with both the Midbrain and the Spinal Cord B region, namely *NPPA* and *UNC5B*. *NPPA* encodes for atrial natriuretic peptide (ANP). In the brain, ANP was described to regulate fluid homeostasis and blood-brain barrier (BBB) permeability. However, ANP was also shown to act as neuromodulator, leading to increased levels of the second messenger cGMP and corresponding downstream pathways [43, 44]. In a similar manner, *UNC5B* was shown to have dual functions. Interactions between UNC5B and netrins are known axon guidance molecules during developmental stages, but the same signalling pathway was shown to regulate blood vessel guidance [45]. This consequently underlines that blood vessel formation is an important feature of tumour infiltration. Interestingly however, netrin-1 was found elevated in medulloblastoma (MB) tumours and netrin-1/UNC5B interactions were shown to promote metastasis in non-subgroup-specific MB [46]. More specifically, interactions between this ligand-receptor pair in MB activate the MAPK pathway, particularly Erk1/2 signalling [47]. The authors further show that UNC5B silencing suppressed MB cell invasion in vitro. This axis may therefore exert similar functions in DIPG.

**Figure 4:**
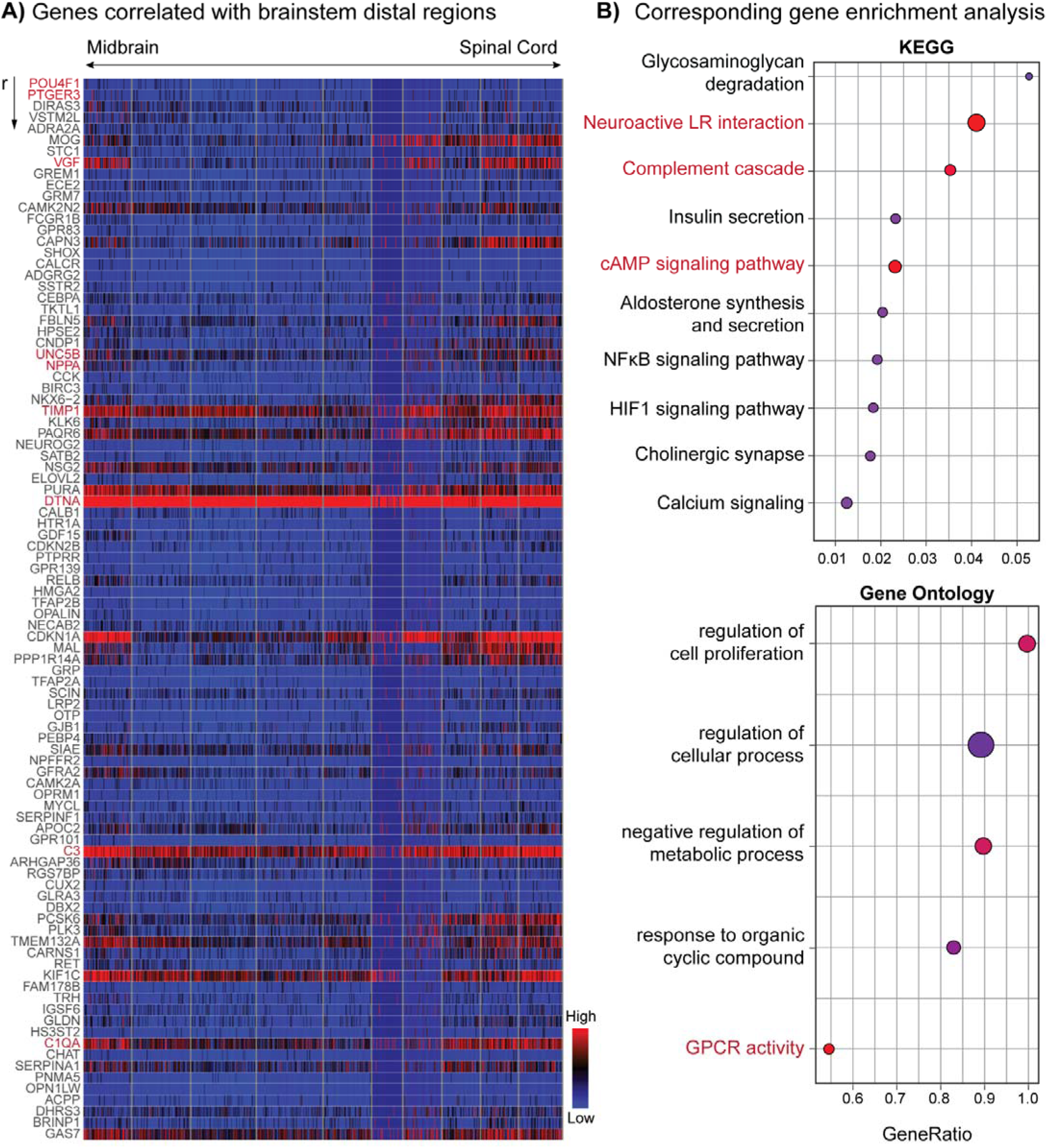
Assessment of gene signatures correlated to the tumour infiltrative edges. **(A)** Heatmap showing 94 genes that are significantly (p>0.049) and positively correlated with tumour distal areas compared to proximal regions. Genes ranking from top to bottom according to Pearson correlation coefficient (r), comparing Pons D to the Midbrain and Spinal Cord D region. **(B)** Gene enrichment analysis of genes identified in (A) demonstrates that the infiltrative edge is most highly associated with neuronal gene signatures and cAMP/GPCR signalling.

Overall, our findings are consistent with neuron-to-glia interactions driving tumour cell invasion and suggest novel targetable candidates for therapeutic intervention.

### DIPG tumour cells interact with endothelial, neuronal and myeloid cell populations

Gene expression profiling in the spatial context offers a significant opportunity to uncover important cell-cell (here community) interactions (CCIs) of a given sample. Information about known ligand-receptor (LR) pairs can be utilised to predict intercellular communication, based on the coordinated expression of genes in particular regions. We used stLearn to assess spatial neighbourhoods of LR co-expressions to compute corresponding LR scores. We were then able to make inferences about cell community interactions (**Figure 5A**). CCI scores were calculated for all ten cell communities and all ten brainstem regions respectively. Whilst certain region-specific differences were observed, the endothelial cell population was the strongest interacting partner in the majority of the ten brainstem areas (**Figure 5B and Supplementary Figure S4**). To further quantify significant CCIs in the DIPG-infiltrated brainstem as a whole, scores of all regions were combined and averaged. Interestingly, we observed clear differences between sender (ligand-expressing) and receiver (receptor-expressing) populations. For instance, the endothelial population was identified as the strongest interacting ligand-expressing community. This was followed by the neuronal and myeloid populations. On the flipside, the two tumour cell communities C1 and C0 were demonstrated to be the most important receiver populations (**Figure 5C**). These results clearly demonstrate that DIPG tumour cells actively receive cues from the TME and underline the importance of cellular communication, particularly between tumour, endothelial, neuronal and myeloid cells. Hence, interrupting such interactions will be an important strategy to combat DIPG.

**Figure 5:**
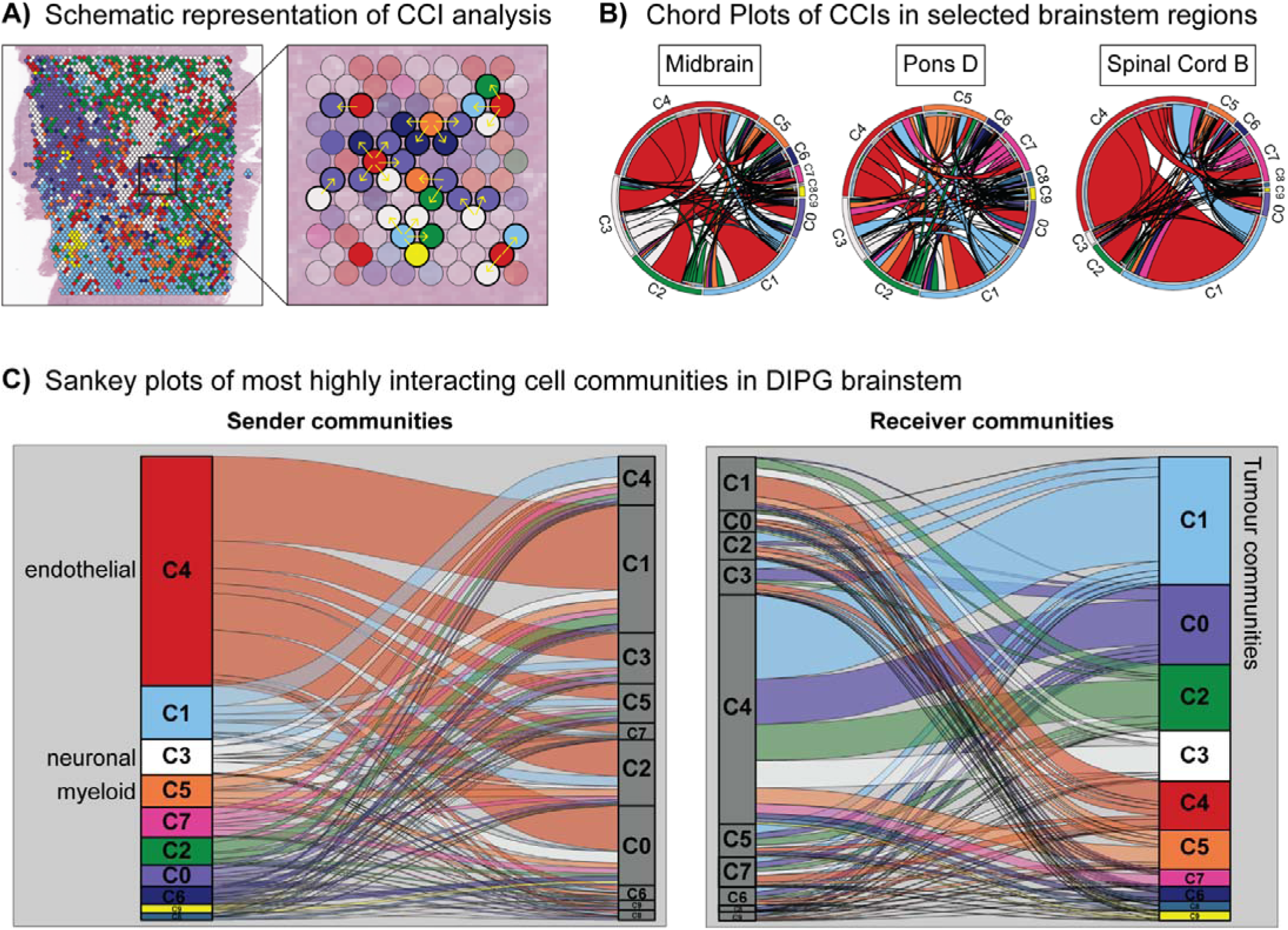
Cell-cell interaction profiling of 10 DIPG cell communities. **(A)** Schematic representation of cell-cell interaction (CCI) analysis. Interactions were predicted based on spot proximity and correlated gene expression. **(B)** Chord plots of 3 exemplary brainstem regions showing the complex interaction network between the 10 DIPG cell communities. Endothelial populations (C4) interact most strongly with other populations. **(C)** Sankey plots visualising cell communities that are most significantly involved in ligand-expressing (sender) or receptor-expressing (receiver)-mediated interactions. Endothelial, neuronal and myeloid communities are amongst the top senders, whilst the tumour populations C1 and C0 are the most significant receiver communities.

### Prediction analysis of ligand-receptor pairs sheds light on novel candidates that contribute to DIPG progression

To evaluate which ligand-receptor pairs were involved in the above described CCIs, we focused on the three most significant interacting partners: endothelial-to-tumour, neuronal-to-tumour and myeloid-to-tumour. Interaction scores were assessed for each brainstem region respectively (**Supplementary Tables S1-S3**). From these, we further determined those interaction partners that a) occur in more than one of the brainstem regions and b) are involved in the communication with more than one tumour cell community. This was to identify those LR pairs that likely play a substantial role for the interaction between DIPG tumour cells and the TME. Focussing on endothelial-to-tumour interactions, *ENG*-*BMP7* (endoglin/bone morphogenic protein7) was identified as the most significant and most frequently recurring LR pair (**Figure 6A**). BMP7 belongs to the transforming growth factor-beta (TGF-β) family and has previously been associated with a glioma quiescent stem cell populations [48]. This is consistent with the vascular niche reflecting a known repository for glioma stem cells [49]. Gliomas further utilise the vascular niche to migrate along blood vessels and most recent studies on BMP7 suggest that signalling through this cell surface receptor promotes DIPG invasion [50]. Expression profiling of endoglin and BMP7 in our dataset confirmed that endoglin is exclusive to the endothelial communities (C4 and C7), whilst *BMP7* is highly associated with tumour cell communities (**Figure 6B**). We also found *TIMP1*-*CD63* among the top 10 LR pairs involved in endothelial-to-glioma interactions. Similar to BMP7 signalling, this pathway is described to maintain a stem-like phenotype in glioma [40]. We further looked at regions in which interactions between the respective LR pairs are predicted to occur and found that the endothelial community C4 interacts most highly with the more immature tumour cell community C1 (**Figure 6C.I and Figure 5C**). This further underlines the vascular niche as a safeguard for the DIPG stem cell pool. Interaction profiling of neuronal-to-tumour communities revealed *ADCYAP1*-*ADCYAP1R1* and *ADCYAP1*-*VIPR2* as the most significant LR pairs (**Figure 6A**). These genes encode for the pituitary adenylate cyclase-activating polypeptide PACAP and the corresponding receptors PAC1 and vasoactive intestinal peptide receptor 2 (VPAC2). PACAP is a neuropeptide, which belongs to the vasoactive intestinal polypeptide (VIP) family and is primarily expressed in nervous tissues where it acts as neurotransmitter and neuromodulator [51]. PAC1 and VPAC2 belong to the GPCR family. Interactions between these ligand-receptor pairs mainly activate the cAMP/PKA (protein kinase A) signalling pathway [52]. Notably, PACAP-PAC1/VPAC2-mediated cAMP/PKA was shown to regulate an invasive phenotype in adult GBM [53]. Additionally, Shioda and colleagues demonstrated that PECAP-PAC1 interactions lead to increased calcium (Ca^2+^) concentrations in rat neuroepithelial cells and downstream activation of the PLC/PKC (phospholipase C/protein kinase C) cascade [54]. Ca^2+^-dependent PLC and PKC signalling is widely implicated in synaptic plasticity and the importance of Ca^2+^ signalling to modulate synaptic strength in DIPG is well established [39, 55, 56]. Most intriguingly, the PACAP-mediated pathway has been implicated to play a notable role in chronic migraine and a phase II clinical trial (NCT04976309) targeting this neuropeptide is currently under investigation [57–59]. Expression analysis of *ADCYAP1*-*ADCYAP1R1* in our DIPG spatial object confirms *ADCYAP1* is restricted to the neuronal community (C3) and *ADCYAP1R1* strongly correlated with tumour cell communities (**Figure 6B**). We also determined regions of *ADCYAP1*-*ADCYAP1R1* interactions in the spatial context. In contrast to endothelial-to-tumour interactions, regions of high neuron-to-glioma communication were predominantly found amongst the more mature tumour population C0 (**Figure 6C.II**). Interestingly, the corresponding H&E of *ADCYAP1*-*ADCYAP1R1*-high areas with an example of the midbrain region shows what appear to be actively infiltrating tumour cells surrounding neurons. This finding is consistent with the body of evidence that neuron-to-glioma interactions crucially mediate glioma progression [16–18, 30, 60]. Hence, it is tempting to speculate that PACAP-PAC1 interactions contribute to neuron-to-glioma-mediated DIPG cell invasion and that blocking PACAP-mediated signalling could provide a strategy to block synaptic communication. Lastly, we found complement component C3 to be greatly involved in immune-to-tumour communication (**Figure 6A**). This is in line with our previous finding, suggesting this candidate contributes to DIPG invasion. Indeed, C3 has been demonstrated to alter a broad range of mitogenic signalling pathways in physiology and pathophysiology, including sustained oncogenesis, invasion and migration, angiogenesis or decreased apoptosis [61–65]. Crosstalk between myeloid cells and glioma cells via C3 were reported previously. CD46 for instance, is considered to facilitate immune escape mechanisms. In line with the previous note, this cell surface receptor was reported to be expressed on glioma stem cells (GSCs) and interactions with C3 shown to support evasion of complement-mediated immune responses [66]. Similarly, a very recent study found that C3a/C3aR pairing between tumour-associated macrophages and GSCs maintains the malignant phenotype of this population [67]. Consistent with these findings, spatial profiling of myeloid-to-tumour-high areas in our dataset displayed myeloid clusters neighbouring the more immature tumour community C1 (**Figure 6C.III**). The same areas also exhibit higher expression of LRP1. This receptor was demonstrated to promote GBM invasion and to be elevated in DIPG [68, 69]. Expression profiling in our dataset confirmed this observation (**Figure 6B**).

**Figure 6:**
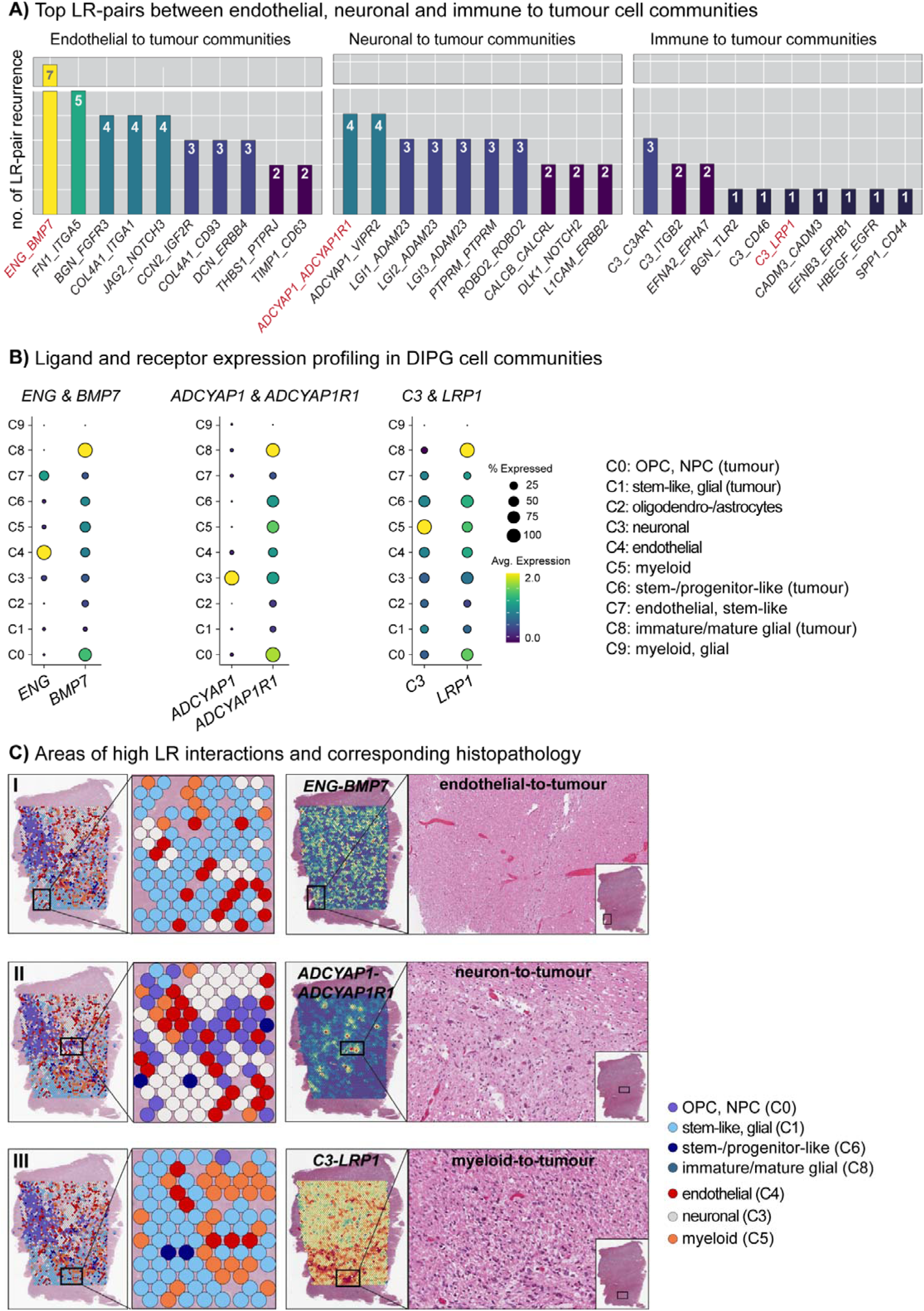
Ligand-receptor pairs that are highly involved in communication between tumour populations and the TME. **(A)** The top 10 ligand-receptor (LR) pairs contributing to interactions between endothelial-to-tumour, neuronal-to-tumour and myeloid-to-tumour were determined. Endoglin (ENG)-BMP7 is the most significant pair and involved in endothelial-to-tumour communications. ADCYAP1 strongly contributes to neuronal-to-tumour interactions and pairs with the corresponding receptor ADCYAP1R1. Complement C3 is the main contributor to myeloid-to-tumour communication. **(B)** Expression profiling of selected LRs confirming that ligands are exclusive to the respective TME interaction partner, whilst the corresponding receptors are strongly associated with tumour cell communities. **(C)** Spatially-informed profiling of predicted LR interactions outlined in (A) together with corresponding H&E images clearly demonstrate patterns of expected pathophysiology. Endothelial and myeloid populations mainly neighbour the more immature tumour cell community C1, neuronal-to-tumour interactions occur mostly with the tumour community correlated to known DIPG markers (C0).

Overall, our LR pair prediction analysis not only highlights the importance of the vascular niche and the contribution of neuronal and myeloid cells to DIPG disease progression, but notably, uncovers targetable candidates involved in these malignant processes. Some of the LR pairs we identified are well known to promote pathophysiology in various diseases and hence, interrupting these communication axes could prove useful to slow DIPG progression.

## DISCUSSION

Many sequencing studies of DIPG tumours have been conducted at the single cell and spatiotemporal level over the past decades. However, complete profiling of an intact tumour-infiltrated brainstem from a DIPG patient has yet to be reported. Our spatial whole transcriptome sequencing study of ten consecutive tumour regions, spanning the entire brainstem from midbrain down to the spinal cord is therefore meaningful and will hopefully add further knowledge to this rapidly progressing field.

Spatially-resolved studies from Ren as well as Filbin and colleagues suggested the presence of region-specific tumour niches that favour modes of tumour cell differentiation and the development of distinct tumour subclones [13, 70]. Our analysis uncovered four genetically diverse tumour cell populations that represent varying states of stemness and maturity, likely reflecting distinct tumour subclones [25, 27, 71]. Spatial trajectory analysis further supported this hypothesis, predicting that these populations transition from immature to more committed tumour cells. These results consequently support the notion that DIPG tumours are comprised of multifaceted tumour niches that give rise to specific tumour phenotypes, which likely contribute functionally to tumour diversity. Thus, our findings and previous studies now lend themselves to further investigation unravelling which molecular mechanisms directly promote such phenotype switches.

Furthermore, our resource allowed us to track tumour invasive signatures and to assess gene expression alterations between distal regions compared to the likely initiation site. We found genes that were predominantly correlated to transcriptional states that reflect neuronal development and signalling. The corresponding effector pathways are mainly defined by GPCR, cAMP and calcium signalling. In concordance with the growing field of cancer neuroscience, our evaluation thereby suggests neuron-mediated cell invasion and determined interesting candidates that could serve as putative treatment targets.

Importantly, the power of spatially-resolved sequencing allowed us to study cellular communication between the heterogeneous cell populations present in this disease, particularly between tumour cells and the TME. As the dominant receiver populations, our CCI analysis highlighted that tumour cells extensively receive cues from neighbouring populations, rather than releasing such cues themselves. Not surprisingly, the vascular niche appears centrally involved in tumour-to-TME interactions, underscoring the importance of the tumour vascular bed for glioma growth and treatment resistance [72]. Aside from this, we also found that neuronal and myeloid cells strongly signal to tumour communities. Interestingly, neurons seem to interact more with tumour cells that are defined by known DIPG markers such as *NTRK2*, *EGFR* or *OLIG2*, whereas endothelial and myeloid populations seem to connect to stem cell-like populations, marked by genes like *FOS* or *JUNB*. Further investigation will be required to confirm this observation, but we speculate that different DIPG subpopulations favour specific processes of disease progression. More importantly however, this analysis enabled us to identify specific ligand-receptor (LR) pairs that are involved in such cellular communication. Our assessment of the top contributing LR pairs uncovered interactions that are largely described to contribute to pathophysiology, but are novel to DIPG malignancy. PACAP-PAC1 of particular note. This LR pair was most highly involved in neuron-to-tumour interactions in our dataset, but more interestingly, phase II clinical trials against both PACAP (NCT04976309) and PAC1 (NCT03238781) are under investigation for migraine prevention [58, 73]. This likely indicates that the respective compounds, both humanised monoclonal antibodies, demonstrate sufficient BBB penetration with low CNS toxicity. However, expression levels of PACAP and PAC1 remain to be validated at the protein level in a cohort of DIPG patient tissues with subsequent *in vitro* and *in vivo* efficacy studies to be conducted in the future. Moreover, the 10x Visium approach lacks single cell resolution. Hence, additional multimodal studies of regionally distinct tumour samples from the same patient should be conducted to further confirm our findings.

Spatially-resolved characterisation of DIPG holds great potential to better understand the underlying disease biology and to ultimately improve patient outcomes. Our study emphasised the importance of immune escape mechanisms, maintenance of a stem cell-like phenotype and neuron-to-glioma interactions in the DIPG setting and, importantly, suggested novel disease contributors that could be addressed therapeutically to impede DIPG tumour progression. Collectively, this study provides useful and relevant information regarding DIPG progression and invasion in its entirety and our novel spatial dataset will hopefully inform future DIPG discovery and translational efforts.

## METHODS

### Patient tissue sampling and ethical consideration

The DIPG tumour donation used for this study was deidentified and obtained through the QCTB with consent of the patient’s legal representative. Sample processing was performed with ethical approval from the QIMR human research ethics committee (P3420). The specimen was collected within hours of the patient’s death and spanned the entire brainstem, including midbrain, pons, medulla oblongata and upper spinal cord regions. Tissue pieces were collected from each region and subsequently formalin-fixed for the preparation of FFPE tissue blocks.

### Tissue fixation and paraffin embedding

Fresh tissues were formalin-fixed for at least 48 hours before the formalin solution was replaced with 70% ethanol (EtOH). Tissues were further cassetted and embedded into paraffin.

### Immunohistochemistry (IHC)

FFPE tissue sections were deparaffinised, rehydrated and tissue epitopes masked during the FFPE process retrieved with a heat-induced method, using a pressure cooker and 10% [w/v] sodium citrate, pH 6.0. Tissues were stained, using the antihistone H3 (mutated K27M) antibody (abcam, ab240310) 1:500 and Hoechst nuclear stain 1:10,000. Images were processed on the Aperio FL Slide Scanner. H&E (hematoxylin and eosin) staining was performed by the QIMRB Histology Core Facility and processed on the Aperio AT Turbo Brightfield Scanner.

### RNA quality control of FFPE tissue blocks

The proportion of RNA fragments with a size of at least 200 nucleotides (DV200) was determined for all tissue samples prior to sequencing. 2x cuts a 10µm per sample were collected and total RNA extracted using the RNeasy FFPE kit (QIAGEN). RNA concentration was determined on the Nanodrop and 50ng/μl used for DV200 indexing employing the Agilent Tapestation.

### Tissue preparation, probe design and library constructions

Spatial transcriptomic sequencing was performed with the 10x Visium CytAssist Spatial Gene Expression platform for FFPE sections. Steps were followed according to the Visium CytAssist for FFPE sections User Guide. In brief, DIPG FFPE tissue blocks were rehydrated under moist conditions in RNAse-free H_2_0 for 1 hour at 4°C prior to sectioning. FFPE blocks were cut at a thickness of 5μm on a microtome. Sections were allowed to fully flatten in a 42°C water bath and transferred onto a microscope glass slide. Slides were placed on a 42°C hot plate for 3 hours and further dried overnight in a glass jar containing desiccator beads. Tissue slides were deparaffinised, stained (H&E) and imaged according to standard histology workflows on day two. Upon imaging and chemical decrosslinking, the slides were placed into a Visium CytAssist Tissue Slide Cassette with a 6.5×6.5mm gasket and prepared for probe hybridisation overnight. Probe ligation, release and extension were performed on the following day. A qPCR step was included to determine the PCR cycle number (C_q_ value) for sample indexing and cDNA amplification. Sample fragment size and sample quality were assessed accordingly for library constructions and sequencing.

### Alignment and quality control (QC)

Raw FASTQ files were processed employing the SpaceRanger software and mapped to the human genome (hg38). Gene expression was quantified based on unique molecular identifier (UMI) counts. For QC adjustment, low quality spots with <2,500 and >75,000 gene counts were removed on each sample individually. The content of mitochondria genes was assessed, but not filtering applied.

### Data integration, spatially-informed clustering and cell type identification

The data of each sample was first merged and further integrated using the anchor-based data integration workflow (nfeatures=3000, normalisation.method=SCTransform) from the Seurat package (v.4.3.0). The *FindClusters* function was used for unsupervised cell type clustering. Based on Clustree (v0.5.0), a resolution of 0.3 was used for clustering. Marker genes for each cell community were determined with the *FindAllMarkers* (only.pos = T, p_val_adj < 0.05) function and cell types identified, using the gene set enrichment analysis tool EnrichR [74].

### CNV estimation for the prediction of tumour content

To identify evidence of large-scale chromosomal copy number variations (CNVs), the InferCNV package (v1.14.1) was utilised. The gene expression data from three human dorsolateral pre-frontal cortex (DLPFC) sections (151507, 151669, 151673), obtained from *spatialLIBD* [75] (https://bioconductor.org/packages/spatialLIBD) were used as a normal reference. The data from the ten DIPG sections and the three normal references was integrated as described above to generate a SeuratObject. With this, a grouped annotation file was provided according to normal or tumour sample (ref_group_names=section). The human genecode hg38 from TrinityCTAT was chosen as a gene order file providing the chromosomal location for each gene. CNV predictions were performed using the six-state Hidden Markov Model (HMM=T) with a specified threshold of 0.2 (BayesMaxPNormal=0.2). The resulting metadata from the InferCNV object was added to the integrated spatialLIBD/DIPG SeuratObject for downstream analysis and plotting.

### Identification of genes associated with brainstem distal regions

To assess genes positively correlated with brainstem distal regions, a matrix with the average expression of all genes (18,025) was generated for each brainstem sample. A correlation vector from either Medulla A to Midbrain or Pons D to Spinal Cord B was assigned and a *cor.test* (method=pearson) performed, determining significance (p) and correlation (r) values. Genes below p<0.049 were removed.

### Spatial trajectory interference, Ligand-Receptor (LR) and Cell-Cell Interaction (CCI) analysis

Analysis of spatial trajectory interference, LR pairing and CCIs was conducted, using the Python-based software stLearn (v.0.4.12) as described previously [76]. Tissue sections from each region were filtered by parameters (min_genes=3, min_cells=100) and then normalised by library size individually. The processed data were subsequently analysed using the stLearn spatial trajectory inference algorithm, which reconstructs cellular trajectories within tissue sections by leveraging spatial distances, morphological features, and gene expression data. Community C6 was defined as the root, and the spatial distribution of communities C0, C1, C6 and C0 trajectories was thoroughly mapped to uncover cellular dynamics and interactions within the tissue context. LR co-expression profiling and equivalent expression levels (LR_SCORE_) between neighbouring spots and between two cell types were based on the repository CellPhoneDB [77]. P-values for each spot and LR pair (p_s,LR_) were used to calculate the CCI_LR_ matrix. CCIs and LR pairs were visualised with RStudio (v.4.2.0), using ggplot and Seurat’s SpatialFeaturePlot function.

## Data and code availability

All data are available from the corresponding author upon request. Source data and codes will be made publically accessible with the final version of the manuscript.

## Author contributions

A.K., Q.N., L.H. and B.D. conceptualisation. A.K. and U.B. processed the autopsy patient tissue. A.K. and T.V. processed the samples for sequencing. A.K., O.M., X.T. and Q.N. analysed the data. B.G.W., T.E.G.H. and M.K.M. contributed intellectually, T.E.G.H. and M.K.M. provided DIPG patient autopsy donation and case report. A.K. wrote the manuscript, L.H. and B.D. review and editing. All authors have reviewed the manuscript.

## Acknowledgements

We thank the QCTB for providing autopsy brain tissue samples and QIMR Histology Core Facility for preparing FFPE tissue blocks. This study was supported by funding from the Sid Faithfull Group and the CHF Children’s Brain Cancer Center.

## REFERENCES

1. Johung, T.B. and M. Monje, Diffuse Intrinsic Pontine Glioma: New Pathophysiological Insights and Emerging Therapeutic Targets. Curr Neuropharmacol, 2017. 15(1): p. 88–97.

2. Warren, K.E., et al., Genomic aberrations in pediatric diffuse intrinsic pontine gliomas. Neuro Oncol, 2012. 14(3): p. 326–32.

3. Monje, M., et al., Hedgehog-responsive candidate cell of origin for diffuse intrinsic pontine glioma. Proceedings of the National Academy of Sciences, 2011. 108(11): p. 4453–4458.

4. Bender, S., et al., Reduced H3K27me3 and DNA hypomethylation are major drivers of gene expression in K27M mutant pediatric high-grade gliomas. Cancer Cell, 2013. 24(5): p. 660–72.

5. Castel, D., et al., Histone H3 wild-type DIPG/DMG overexpressing EZHIP extend the spectrum diffuse midline gliomas with PRC2 inhibition beyond H3-K27M mutation. Acta Neuropathologica, 2020. 139(6): p. 1109–1113.

6. Wu, G., et al., Somatic histone H3 alterations in pediatric diffuse intrinsic pontine gliomas and non-brainstem glioblastomas. Nat Genet, 2012. 44(3): p. 251–3.

7. Duchatel, R.J., et al., Signal Transduction in Diffuse Intrinsic Pontine Glioma. PROTEOMICS, 2019. 19(21-22): p. 1800479.

8. Khuong-Quang, D.-A., et al., K27M mutation in histone H3.3 defines clinically and biologically distinct subgroups of pediatric diffuse intrinsic pontine gliomas. Acta Neuropathologica, 2012. 124(3): p. 439–447.

9. Lapin, D.H., M. Tsoli, and D.S. Ziegler, Genomic Insights into Diffuse Intrinsic Pontine Glioma. Frontiers in Oncology, 2017. 7.

10. Mackay, A., et al., Integrated Molecular Meta-Analysis of 1,000 Pediatric High-Grade and Diffuse Intrinsic Pontine Glioma. Cancer Cell, 2017. 32(4): p. 520–537 e5.

11. Louis, D.N., et al., *The* 2021 WHO Classification of Tumors of the Central Nervous System: a summary. Neuro-Oncology, 2021. 23(8): p. 1231–1251.

12. Filbin, M.G., et al., Developmental and oncogenic programs in H3K27M gliomas dissected by single-cell RNA-seq. Science, 2018. 360(6386): p. 331–335.

13. Liu, I., et al., The landscape of tumor cell states and spatial organization in H3-K27M mutant diffuse midline glioma across age and location. Nature Genetics, 2022. 54(12): p. 1881–1894.

14. Barron, T., et al., GABAergic neuron-to-glioma synapses in diffuse midline gliomas. bioRxiv, 2022: p. 2022.11.08.515720.

15. Taylor, K.R., et al., Glioma synapses recruit mechanisms of adaptive plasticity. bioRxiv, 2021: p. 2021.11.04.467325.

16. Venkataramani, V., et al., Glutamatergic synaptic input to glioma cells drives brain tumour progression. Nature, 2019. 573(7775): p. 532–538.

17. Venkatesh, H.S., et al., Neuronal Activity Promotes Glioma Growth through Neuroligin-3 Secretion. Cell, 2015. 161(4): p. 803–16.

18. Venkatesh, H.S., et al., Electrical and synaptic integration of glioma into neural circuits. Nature, 2019. 573(7775): p. 539–545.

19. Venkataramani, V., et al., Disconnecting multicellular networks in brain tumours. Nat Rev Cancer, 2022. 22(8): p. 481–491.

20. Long, Y., et al., Spatially informed clustering, integration, and deconvolution of spatial transcriptomics with GraphST. Nature Communications, 2023. 14(1): p. 1155.

21. Erickson, A., et al., Spatially resolved clonal copy number alterations in benign and malignant tissue. Nature, 2022. 608(7922): p. 360–367.

22. Maynard, K.R., et al., Transcriptome-scale spatial gene expression in the human dorsolateral prefrontal cortex. Nat Neurosci, 2021. 24(3): p. 425–436.

23. Warren, K.E., et al., Genomic aberrations in pediatric diffuse intrinsic pontine gliomas. Neuro-Oncology, 2011. 14(3): p. 326–332.

24. Hoffman, L.M., et al., Spatial genomic heterogeneity in diffuse intrinsic pontine and midline high-grade glioma: implications for diagnostic biopsy and targeted therapeutics. Acta Neuropathol Commun, 2016. 4: p. 1.

25. Nikbakht, H., et al., Spatial and temporal homogeneity of driver mutations in diffuse intrinsic pontine glioma. Nat Commun, 2016. 7: p. 11185.

26. Tari, H., et al., Quantification of spatial subclonal interactions enhancing the invasive phenotype of pediatric glioma. Cell Reports, 2022. 40(9): p. 111283.

27. Vinci, M., et al., Functional diversity and cooperativity between subclonal populations of pediatric glioblastoma and diffuse intrinsic pontine glioma cells. Nat Med, 2018. 24(8): p. 1204–1215.

28. Krishna, S., et al., Glioblastoma remodelling of human neural circuits decreases survival. Nature, 2023. 617(7961): p. 599–607.

29. Monje, M., et al., Roadmap for the Emerging Field of Cancer Neuroscience. Cell, 2020. 181(2): p. 219–222.

30. Venkataramani, V., et al., Glioblastoma hijacks neuronal mechanisms for brain invasion. Cell, 2022. 185(16): p. 2899–2917.e31.

31. Venkatesh, H.S., et al., Targeting neuronal activity-regulated neuroligin-3 dependency in high-grade glioma. Nature, 2017. 549(7673): p. 533–537.

32. Badea, T.C., et al., Combinatorial Expression of Brn3 Transcription Factors in Somatosensory Neurons: Genetic and Morphologic Analysis. The Journal of Neuroscience, 2012. 32(3): p. 995–1007.

33. Serrano-Saiz, E., et al., BRN3-type POU Homeobox Genes Maintain the Identity of Mature Postmitotic Neurons in Nematodes and Mice. Curr Biol, 2018. 28(17): p. 2813–2823.e2.

34. Cantaut-Belarif, Y., et al., Microglia control the glycinergic but not the GABAergic synapses via prostaglandin E2 in the spinal cord. J Cell Biol, 2017. 216(9): p. 2979–2989.

35. Schafer, D.P., et al., Microglia sculpt postnatal neural circuits in an activity and complement-dependent manner. Neuron, 2012. 74(4): p. 691–705.

36. Stevens, B., et al., The classical complement cascade mediates CNS synapse elimination. Cell, 2007. 131(6): p. 1164–1178.

37. Alder, J., et al., Brain-derived neurotrophic factor-induced gene expression reveals novel actions of VGF in hippocampal synaptic plasticity. J Neurosci, 2003. 23(34): p. 10800–8.

38. Lin, W.-J., et al., VGF and Its C-Terminal Peptide TLQP-62 Regulate Memory Formation in Hippocampus via a BDNF-TrkB-Dependent Mechanism. The Journal of Neuroscience, 2015. 35(28): p. 10343–10356.

39. Taylor, K.R., et al., Glioma synapses recruit mechanisms of adaptive plasticity. Nature, 2023. 623(7986): p. 366–374.

40. Aaberg-Jessen, C., et al., Co-expression of TIMP-1 and its cell surface binding partner CD63 in glioblastomas. BMC Cancer, 2018. 18(1): p. 270.

41. Blake, D.J., et al., Different Dystrophin-like Complexes Are Expressed in Neurons and Glia. Journal of Cell Biology, 1999. 147(3): p. 645–658.

42. Montanaro, F. and S. Carbonetto, Targeting Dystroglycan in the Brain. Neuron, 2003. 37(2): p. 193–196.

43. de Vente, J., J.G.J.M. Bol, and H.W.M. Steinbusch, cGMP-Producing, Atrial Natriuretic Factor-Responding Cells in the Rat Brain. European Journal of Neuroscience, 1989. 1(5): p. 436–460.

44. Whitson, P.A., M.H. Huls, and C.F. Sams, Characterization of atrial natriuretic peptide receptors in brain microvessel endothelial cells. Journal of Cellular Physiology, 1991. 146(1): p. 43–51.

45. Carmeliet, P. and M. Tessier-Lavigne, Common mechanisms of nerve and blood vessel wiring. Nature, 2005. 436(7048): p. 193–200.

46. Li, M., Y. Deng, and W. Zhang, Molecular Determinants of Medulloblastoma Metastasis and Leptomeningeal Dissemination. Molecular Cancer Research, 2021. 19(5): p. 743–752.

47. Akino, T., et al., Netrin-1 Promotes Medulloblastoma Cell Invasiveness and Angiogenesis, and Demonstrates Elevated Expression in Tumor Tissue and Urine of Patients with Pediatric Medulloblastoma. Cancer Research, 2014. 74(14): p. 3716–3726.

48. Sachdeva, R., et al., BMP signaling mediates glioma stem cell quiescence and confers treatment resistance in glioblastoma. Scientific Reports, 2019. 9(1): p. 14569.

49. Gilbertson, R.J. and J.N. Rich, Making a tumour’s bed: glioblastoma stem cells and the vascular niche. Nature Reviews Cancer, 2007. 7(10): p. 733–736.

50. Bruschi, M., et al., Abstract 154: Diffuse midline glioma invasion and metastasis rely on cell-autonomous signaling through a phenotypic switch controlled by BMP7. Cancer Research, 2024. 84(6_Supplement): p. 154–154.

51. Shioda, S., Pituitary adenylate cyclase-activating polypeptide (PACAP) and its receptors in the brain. Kaibogaku Zasshi, 2000. 75(6): p. 487–507.

52. Hirabayashi, T., T. Nakamachi, and S. Shioda, Discovery of PACAP and its receptors in the brain. The Journal of Headache and Pain, 2018. 19(1): p. 28.

53. Bensalma, S., et al., PKA at a Cross-Road of Signaling Pathways Involved in the Regulation of Glioblastoma Migration and Invasion by the Neuropeptides VIP and PACAP. Cancers, 2019. 11(1): p. 123.

54. Zhou, C.-J., et al., PACAP activates PKA, PKC and Ca2+ signaling cascades in rat neuroepithelial cells. Peptides, 2001. 22(7): p. 1111–1117.

55. Hongpaisan, J. and D.L. Alkon, A structural basis for enhancement of long-term associative memory in single dendritic spines regulated by PKC. Proceedings of the National Academy of Sciences, 2007. 104(49): p. 19571–19576.

56. Olds, J.L., et al., Imaging of Memory-Specific Changes in the Distribution of Protein Kinase C in the Hippocampus. Science, 1989. 245(4920): p. 866–869.

57. Birk, S., et al., The effect of intravenous PACAP38 on cerebral hemodynamics in healthy volunteers. Regulatory Peptides, 2007. 140(3): p. 185–191.

58. Rasmussen, N.B., et al., The effect of Lu AG09222 on PACAP38- and VIP-induced vasodilation, heart rate increase, and headache in healthy subjects: an interventional, randomized, double-blind, parallel-group, placebo-controlled study. The Journal of Headache and Pain, 2023. 24(1): p. 60.

59. Schytz, H.W., et al., PACAP38 induces migraine-like attacks in patients with migraine without aura. Brain, 2008. 132(1): p. 16–25.

60. Pan, Y., et al., NF1 mutation drives neuronal activity-dependent initiation of optic glioma. Nature, 2021. 594(7862): p. 277–282.

61. Markiewski, M.M., et al., Modulation of the antitumor immune response by complement. Nat Immunol, 2008. 9(11): p. 1225–35.

62. Nozaki, M., et al., Drusen complement components C3a and C5a promote choroidal neovascularization. Proc Natl Acad Sci U S A, 2006. 103(7): p. 2328–33.

63. Rutkowski, M.J., et al., Cancer and the Complement Cascade. Molecular Cancer Research, 2010. 8(11): p. 1453–1465.

64. Tang, Z., et al., C3a mediates epithelial-to-mesenchymal transition in proteinuric nephropathy. J Am Soc Nephrol, 2009. 20(3): p. 593–603.

65. van Beek, J., et al., Complement anaphylatoxin C3a is selectively protective against NMDA-induced neuronal cell death. Neuroreport, 2001. 12(2): p. 289–93.

66. Yuan, D., et al., Systematic expression analysis of ligand-receptor pairs reveals important cell-to-cell interactions inside glioma. Cell Communication and Signaling, 2019. 17(1): p. 48.

67. Zhang, Y., et al., An NFAT1-C3a-C3aR Positive Feedback Loop in Tumor-Associated Macrophages Promotes a Glioma Stem Cell Malignant Phenotype. Cancer Immunology Research, 2024. 12(3): p. 363–376.

68. Karki, A., et al., Receptor-driven invasion profiles in diffuse intrinsic pontine glioma. Neuro-Oncology Advances, 2021. 3(1).

69. Song, H., et al., Low-Density Lipoprotein Receptor-Related Protein 1 Promotes Cancer Cell Migration and Invasion by Inducing the Expression of Matrix Metalloproteinases 2 and 9. Cancer Research, 2009. 69(3): p. 879–886.

70. Ren, Y., et al., Spatial transcriptomics reveals niche-specific enrichment and vulnerabilities of radial glial stem-like cells in malignant gliomas. Nature Communications, 2023. 14(1): p. 1028.

71. Ryall, S.T., et al., Abstract 1184: Clonal evolution of diffuse intrinsic pontine glioma. Cancer Research, 2018. 78(13_Supplement): p. 1184–1184.

72. Jung, E., et al., Tumor cell plasticity, heterogeneity, and resistance in crucial microenvironmental niches in glioma. Nature Communications, 2021. 12(1): p. 1014.

73. Ashina, M., et al., A phase 2, randomized, double-blind, placebo-controlled trial of AMG 301, a pituitary adenylate cyclase-activating polypeptide PAC1 receptor monoclonal antibody for migraine prevention. Cephalalgia, 2021. 41(1): p. 33–44.

74. Kuleshov, M.V., et al., Enrichr: a comprehensive gene set enrichment analysis web server 2016 update. Nucleic Acids Research, 2016. 44(W1): p. W90–W97.

75. Pardo, B., et al., spatialLIBD: an R/Bioconductor package to visualize spatially-resolved transcriptomics data. BMC Genomics, 2022. 23(1): p. 434.

76. Pham, D., et al., Robust mapping of spatiotemporal trajectories and cell–cell interactions in healthy and diseased tissues. Nature Communications, 2023. 14(1): p. 7739.

77. Efremova, M., et al., CellPhoneDB: inferring cell–cell communication from combined expression of multi-subunit ligand–receptor complexes. Nature Protocols, 2020. 15(4): p. 1484–1506.

