## Supplementary Data for "Spatial Transcriptomic Sequencing of a DIPG-infiltrated Brainstem reveals Key Invasion Markers and Novel Ligand-Receptor Pairs contributing to Tumour to TME Crosstalk"

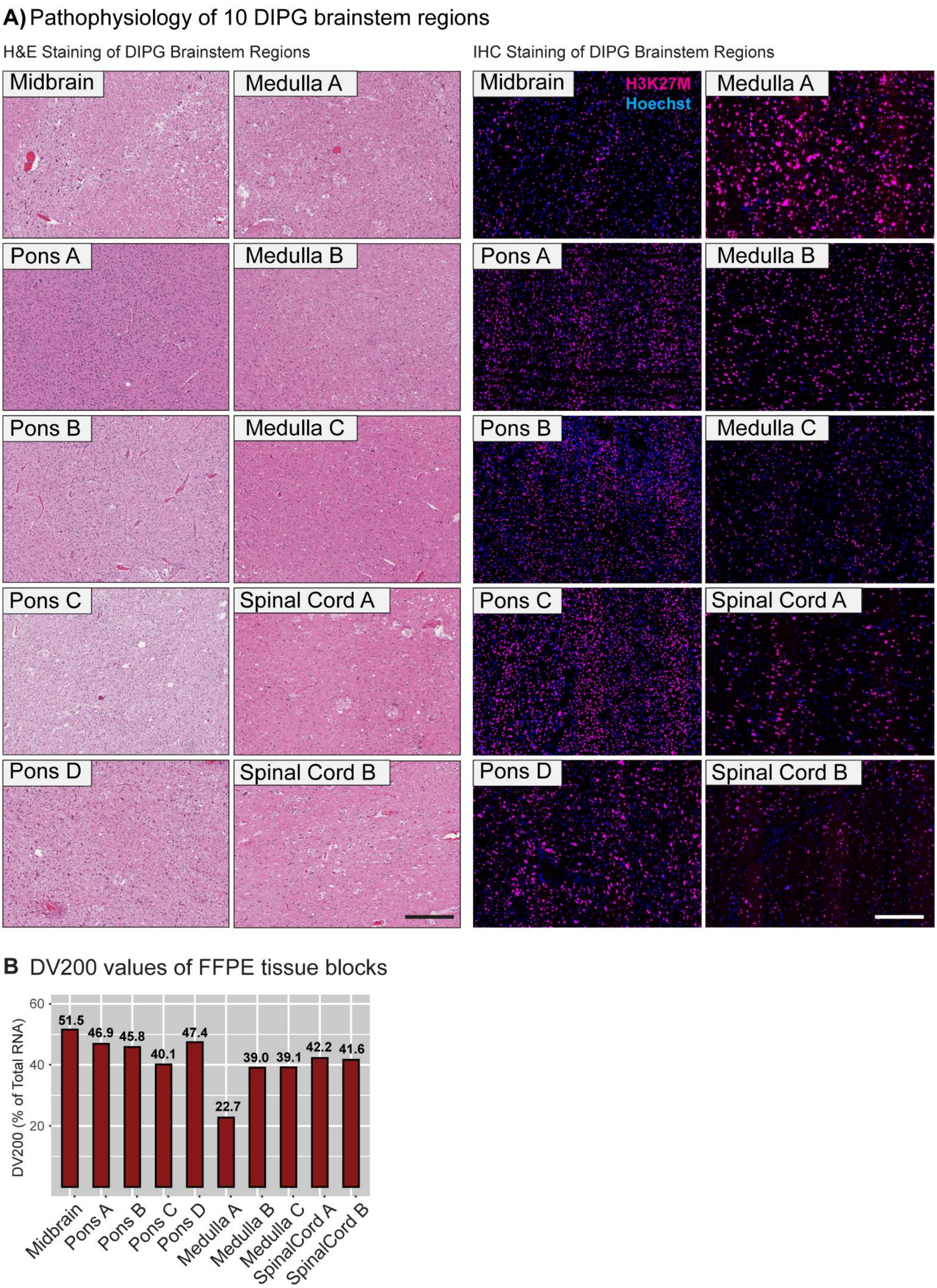

**Supplementary Figure S1: Tumour infiltration patterns across 10 DIPG brainstem regions. (A)** H&E and H3K27M staining confirming DIPG burden in all regions to varying degrees. **(B)** Assessment of FFPE tissue quality demonstrating sufficient RNA quality for sequencing studies. Region Medulla A, which presents areas of tumour necrosis as per staining demonstrated in (A) shows lowest RNA quality. Scale bar is 300µm.

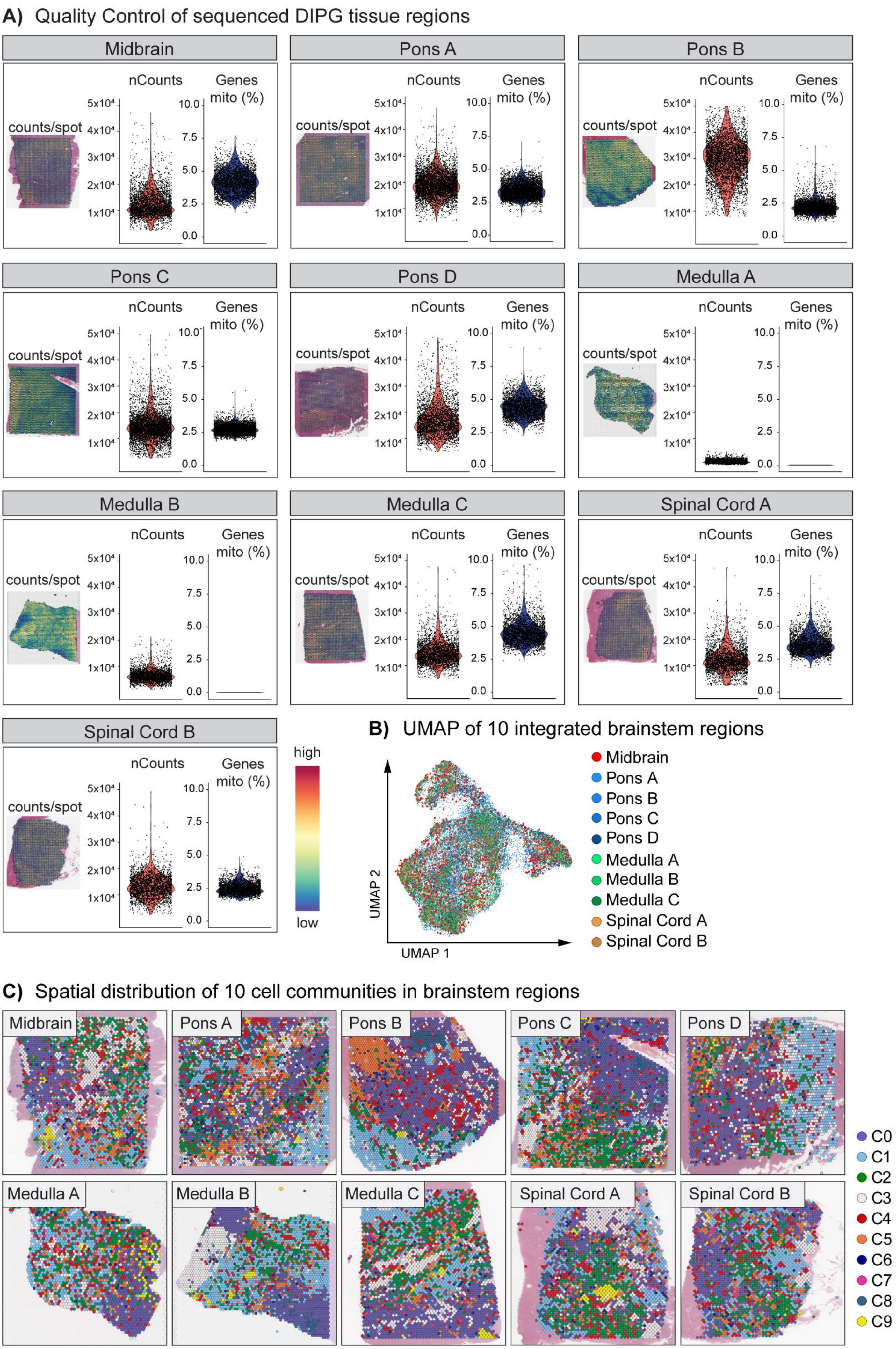
A) Quality Control of sequenced DIPG tissue regions
Midbrain
Pons A
Pons B
Pons C
Pons D
Medulla A
Medulla B
Medulla C
Spinal Cord A
Spinal Cord B
counts/spot
nCounts
Genes mito (%)
5x10⁴
4x10⁴
3x10⁴
2x10⁴
1x10⁴
10.0
7.5
5.0
2.5
0.0
high
low
B) UMAP of 10 integrated brainstem regions
UMAP 2
UMAP 1
Midbrain
Pons A
Pons B
Pons C
Pons D
Medulla A
Medulla B
Medulla C
Spinal Cord A
Spinal Cord B
C) Spatial distribution of 10 cell communities in brainstem regions
Midbrain
Pons A
Pons B
Pons C
Pons D
Medulla A
Medulla B
Medulla C
Spinal Cord A
Spinal Cord B
C0
C1
C2
C3
C4
C5
C6
C7
C8
C9

**Supplementary Figure S2: Quality Control and data integration of sequenced DIPG tumour tissue regions. (A)** The median counts and the mitochondrial gene content was assessed for each of the 10 sequenced DIPG tissue regions. Median counts ranging between 1,420 (Medulla A) and 30,695 (Pons B). The mitochondrial gene content was low for all tissue regions. **(B)** UMAP plotting showing dimensional reduction of the 10 DIPG brainstem regions after merging and anchor-based data integration. **(C)** Spatial visualisation of the 10 cell communities in the 10 DIPG brainstem regions highlighting cellular heterogeneity present in the disease.

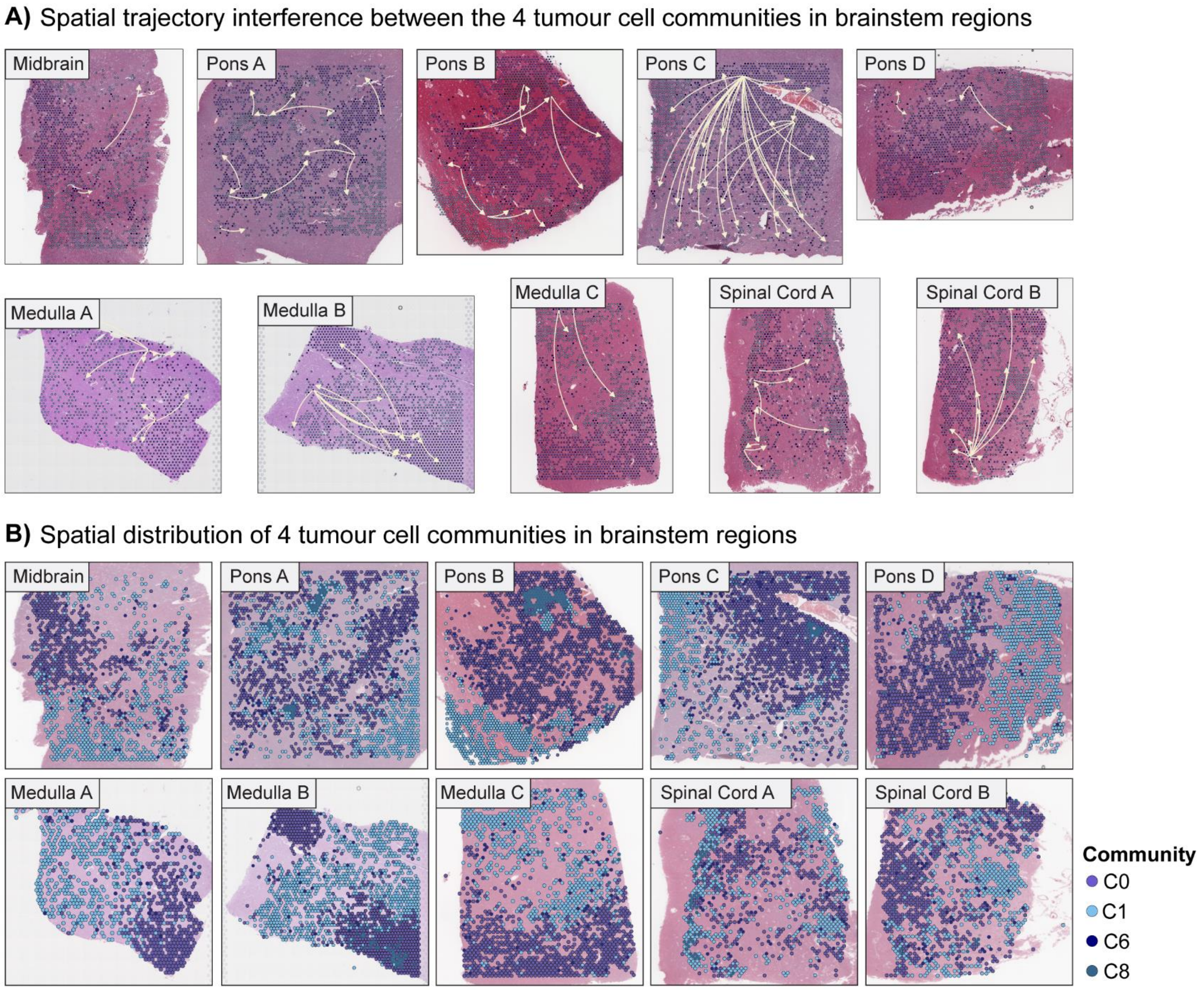

**Supplementary Figure S3: Spatial distribution and trajectory analysis of 4 tumour cell communities in 10 DIPG brainstem regions. (B)** Visualisation of the 4 tumour-associated cell populations shows different degrees of compartmentalisation. C0 and C8 appear to cluster most strongly with themselves, whereas C6 is widely dispersed. C1 largely appears clustered but more widely distributed across the tissue compared to C0 and C8. **(A)** Pseudo-space-time distance analysis revealing spatial trajectory patterns of the 4 tumour communities, suggestion development form immature to more committed subpopulations (C6 to C1 to C8 to C0).

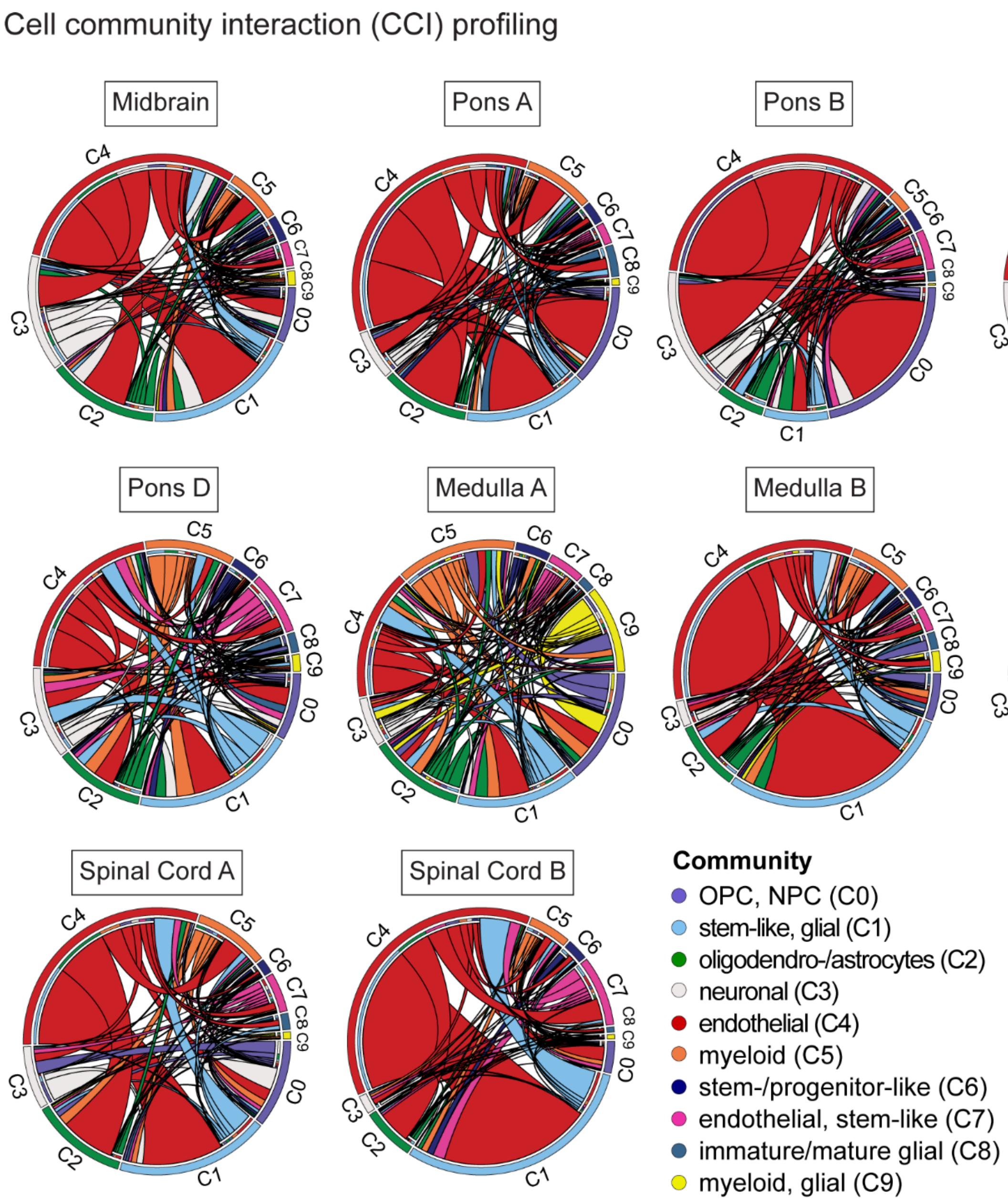

**Supplementary Figure S4: Cell community interaction profiling.** Chord plots showing predicted interactions between the 10 different cell communities for all 10 brainstem regions respectively. The endothelial community (C4) interacts most significantly with other populations, consistently across all 10 samples.

**Supplementary Table S1: Ligand-Receptor Pairs, Endothelial-to-Tumour Communities**

| Midbrain | PonsA | PonsB | PonsC | PonsD | MedullaA | MedullaB | MedullaC | SpinalCordA | SpinalCordB |
|---|---|---|---|---|---|---|---|---|---|
| ANOS1_SDC2 | SEMA7A_ITGB1 | SEMA7A_ITGB1 | FN1_ITGA4 | NTN4_UNC5A | NLGN1_NRXN1 | PDGFB_PDGFRB | PSEN1_NCSTN | JAG1_NOTCH1 | JAG1_NOTCH1 |
| FN1_ITGA4 | FN1_ITGA9 | FN1_ITGA9 | NECTIN2_NECTIN2 | BMP2_ENG | AGT_LRP2 | C1QA_CD93 | WNT5A_MCAM | IL33_IL1RL1 | NECTIN2_NECTIN2 |
| BMP2_ENG | JAG2_NOTCH3 | TGFB3_ENG | FN1_ITGA9 | PTPRM_PTPRM | WNT5A_FZD2 | TGFB1_TGFBR3 | TGFB3_ENG | THBS2_ITGA6 | FN1_ITGA9 |
| EGF_ERBB3 | FSTL1_DIP2A | SEMA3C_PLXND1 | CCN2_IGF2R | WNT5A_LRP5 | EFNA3_EPHA5 | COL4A1_CD93 | FGF17_FGFR3 | JAG1_CD46 | TGFB3_ENG |
| FSTL1_DIP2A | PSEN1_NOTCH3 | JAG2_NOTCH3 | FAT4_DCHS1 | IFNB1_IFNAR2 | AGRN_ATP1A3 | BSG_SLC16A1 | WNT7B_FZD5 | LAMA5_BCAM | BMP2_ENG |
| THBS1_PTPRJ | COL4A1_ITGAV | EFNB2_EPHB2 | SEMA4A_PLXNB1 | AFDN_EPHB3 | COL4A1_ITGAV | ANGPT2_TIE1 | CADM3_CADM3 | COL4A1_ITGAV | DLL3_NOTCH3 |
| AFDN_EPHB3 | PGF_FLT1 | DCN_ERBB4 | PDGFA_PDGFRA | APELA_APLNR | LGI3_ADAM23 | ANOS1_FGFR1 | CCL28_CCR10 | TFPI_VLDLR | TGFB2_ENG |
| SOST_LRP5 | MFGE8_PDGFRB | FSTL1_DIP2A | FSTL1_DIP2A | CD38_PECAM1 | AGRN_DAG1 | DCN_ERBB4 | DLK1_NOTCH3 | PTN_SDC3 | MDK_ITGB1 |
| AFDN_NECTIN3 | SEMA3A_NRP2 | PSEN1_NOTCH3 | AFDN_NECTIN1 | WNT5A_FZD4 | NCAM1_ROBO3 | CCN2_LRP1 | DCN_TLR4 | PGF_FLT1 | JAG1_CD46 |
| TRH_TRHR | FN1_ITGB7 | MFGE8_ITGB3 | APOE_LDLR | PGF_FLT1 | EFEMP1_EGFR | FN1_ITGA5 | IL26_IL20RA | CX3CL1_ITGB3 | LAMA5_BCAM |
| CXCL12_ACKR3 | FN1_CD44 | TGFB3_ACVRL1 | AFDN_NECTIN3 | FN1_ITGB1 | BMP7_BMPR1B | GNAI2_IGF1R | DCN_ERBB4 | NPY_PRLHR | TNFSF15_TNFRSF6B |
| BMP7_ENG | PECAM1_CD38 | MFGE8_PDGFRB | GAS6_AXL | LPL_VLDLR | DLL3_NOTCH2 | FN1_ITGA9 | COL8A1_ITGA1 | FN1_SDC2 | PDGFB_PDGFRB |
| TNFSF13_TNFRSF1A | FN1_ITGA5 | MMP9_EPHB2 | COL3A1_DDR1 | ADAM23_ITGA4 | PTN_PTPRB | CD44_SELE | CXCL10_SDC4 | COL4A1_ITGA1 | TGFB1_ENG |
| ADAM2_ITGA6 | COL1A1_ITGA2 | CXCL12_ITGB1 | VWF_ITGA2B | BGN_TLR2 | LRRC4B_PTPRF | TGFB3_ENG | INHBA_ENG | SEMA3A_PLXNA2 | APOD_LEPR |
| IFNA2_IFNAR2 | SEMA3A_NRP1 | DCN_TLR4 | IFNB1_IFNAR1 | CD160_TNFRSF14 | FGF1_FGFR2 | TGFB2_ENG | IGFBP4_FZD8 | IL18_IL18R1 | PDGFC_PDGFRB |
| FN1_ITGB1 | FN1_TSHR | BMP4_BMPR2 | APOE_LRP8 | BGN_FGFR3 | TGM2_ITGB1 | OSM_IL6ST | DLL3_NOTCH3 | NID1_ITGB3 | JAG1_NOTCH3 |
| BGN_TLR2 | COL1A2_ITGA11 | CCN4_ITGB1 | PGF_FLT1 | COL18A1_ITGB1 | TGM2_SDC4 | DAG1_LAMA4 | TGFB3_ENG | BSG_SELE | FGF2_SDC2 |
| FN1_SDC2 | IGFBP4_FZD8 | NID1_ITGB3 | APOB_LDLR | BSG_SLC16A1 | EFNB3_EPHB3 | PSEN1_NOTCH3 | TGFB2_ENG | SELPLG_SELE | COL18A1_KDR |
| CADM1_CADM1 | FN1_ITGA4 | FN1_ITGA5 | VEGFB_FLT1 | VCAN_TLR2 | GNAS_ADRB2 | COL4A1_ITGAV | JAG2_NOTCH3 | PDGFB_ART1 | PDGFD_PDGFRB |
| EFNB2_EPHB3 | LAMB1_ITGA6 | COL1A1_ITGA2 | COL1A2_ITGB1 | TNFRSF14_CD160 | GNAI2_ADORA1 | GAS6_AXL | COL4A1_ITGAV | BMP6_BMPR2 | VWF_TNFRSF11B |
| CCL8_ACKR1 | BMP2_ENG | THBS1_CD36 | HRAS_INSR | IL4_IL13RA2 | PTN_PTPRZ1 | MFGE8_ITGB3 | MFGE8_ITGB3 | VEGFD_FLT4 | FN1_ITGA9 |
| ICAM2_ITGAL | CD38_PECAM1 | SEMA3A_NRP1 | FNDC5_ITGAV | FN1_ITGA5 | IL1RAPL1_PTPRF | TGFB3_ACVRL1 | LTBP1_ITGB5 | TNC_ITGB3 | TGFB3_ENG |
| THBS2_NOTCH3 | FN1_ITGB3 | CCN2_LRP6 | SEMA6A_PLXNA2 | WNT8A_FZD5 | BMP2_BMPR1B | GNAI2_ADCY1 | NTF3_NGFR | CCN2_ITGB2 | ADAM15_ITGB1 |
| THBS1_ITGA6 | COL1A2_ITGB3 | BMP7_BMPR2 | MATN1_ITGA1 | BGN_TLR4 | VCAN_EGFR | MFGE8_PDGFRB | ANXA1_EGFR | FN1_PLAUR | TGFB2_ENG |
| BGN_FGFR3 | FN1_SDC2 | NTN4_DCC | C3_CD19 | COL4A1_ITGB1 | PTN_PTPRS | LTBP1_ITGB5 | COL1A1_CD93 | COL8A1_ITGA1 | C3_C3AR1 |
| COL1A2_CD93 | A2M_LRP1 | NTNG1_LRRC4C | BMP8A_BMPR2 | WNT5A_ROR1 | HSP90AA1_EGFR | C5_C5AR1 | HBEGF_EGFR | ADAM23_ITGB3 | EFNB2_EPHB2 |
| COL4A1_CD93 | NAMPT_INSR | COL1A2_ITGA11 | CXCL10_CCR3 | BMP2_BMPR1A | CNTN1_NOTCH1 | S100A8_TLR4 | LPL_LRP1 | FN1_ITGA9 | FBLN1_ITGB1 |
| ROBO1_ROBO1 | ANXA1_FPR1 | DLL3_NOTCH3 | SCGB1A1_LRP2 | COL4A1_ITGA1 | TGM2_ADGRG1 | SLITRK5_PTPRD | PECAM1_CD38 | TGFB3_ENG | DCN_ERBB4 |
| DKK2_LRP5 | TGM2_ITGB1 | FN1_ITGA4 | WNT1_FZD3 | THBS1_ITGB1 | COL1A1_DDR1 | APOD_LEPR | FN1_ITGA2B | ADAM15_ITGB1 | PSEN1_NOTCH3 |
| AFDN_NECTIN2 | COL1A1_ITGB1 | OMG_RTN4R | BSG_SLC16A1 | ADM_CALCRL | MFNG_NOTCH2 | NID1_ITGB3 | NPPA_NPR2 | FBLN1_ITGB1 | COL4A1_ITGAV |
| ICAM2_ITGAM | COL18A1_ITGB1 | IHH_BOC | AFDN_NECTIN2 | NECTIN2_NECTIN2 | COL1A1_CD44 | SERPING1_SELE | FN1_ITGA5 | DCN_ERBB4 | FSTL1_DIP2A |
| THBS1_SDC4 | BGN_FGFR3 | TGFB1_ACVRL1 | TGFB1_TGFBR2 | CXCL14_CXCR4 | NCAM1_GFRA1 | INHBA_ACVR1 | FGF5_FGFR2 | PSEN1_NOTCH3 | TNFSF13_TNFRSF14 |

| | | | | | | | | | |
|---|---|---|---|---|---|---|---|---|---|
| GNAI2_OPRM1 | FGF4_NRP1 | LAMB1_ITGA6 | FN1_COL13A1 | FN1_ITGA9 | SLIT2_GPC1 | SELE_CD44 | VCAN_EGFR | COL4A1_ITGAV | ICAM3_CD209 |
| CRH_CRHR2 | BSG_SLC16A1 | IGFBP4_LRP6 | FGF13_SCN5A | FN1_ITGAV | ANOS1_SDC2 | FN1_ITGA5 | LPL_GPIHBP1 | EFNB3_EPHB1 | CD209_ICAM3 |
| PDPN_CLEC1B | SERPINF1_PLXDC1 | TNFSF15_TNFRSF6B | GDF9_BMPR1B | CCN2_IGF2R | C1QB_LRP1 | COL1A1_ITGA2 | EGF_EGFR | DAG1_LAMA4 | PGF_FLT1 |
| FN1_ITGA5 | MDK_ALK | COL3A1_DDR1 | CCN2_ITGB2 | IL33_IL1RL1 | FGF1_FGFR3 | POSTN_PTK7 | L1CAM_EGFR | TSLP_IL7R | MFGE8_PDGFRB |
| THBS1_LRP1 | FN1_ITGA8 | PDGFB_PDGFRB | PDGFD_PDGFRB | DCN_MET | TGFB3_ENG | CCL19_CCRL2 | FN1_ITGA4 | MFGE8_ITGB3 | CXCL12_ITGB1 |
| FN1_ITGA8 | FN1_PLAUR | AHSG_INSR | APOE_SCARB1 | SEMA3B_NRP1 | C3_ITGB2 | CCN2_LRP6 | TGFB1_SDC2 | INHBA_TGFBR3 | JAM2_JAM3 |
| PTPRK_PTPRK | NECTIN1_NECTIN1 | THBS1_ITGA4 | COL3A1_DDR2 | MMRN2_CD93 | B2M_LILRB1 | JAG1_NOTCH4 | IGFBP4_LRP6 | CXCL12_ITGB1 | DCN_TLR4 |
| COL18A1_KDR | ADAM17_ITGA5 | FN1_ITGB3 | SEMA4A_PLXND1 | CXCL5_ACKR1 | TGFB2_ENG | CCN1_ITGB3 | TNFSF12_CD163 | DCN_TLR4 | FN1_ITGB7 |
| THBS1_CD47 | SEMA3B_NRP1 | VEGFB_FLT1 | EFNA5_EPHA1 | CCN2_ITGA5 | LAMA1_ITGB8 | CFH_ITGAM | LGALS3BP_ITGB1 | COL1A1_CD93 | COL1A1_CD93 |
| CXCL12_CD4 | THBS2_ITGA6 | COL1A2_ITGB3 | C3_NRP1 | ADM2_RAMP2 | TGFB1_ITGB8 | GAS6_MERTK | PDGFB_PDGFRB | CD2_CD48 | OSM_OSMR |
| CCL19_CCR10 | CCN2_ITGA5 | FN1_SDC2 | FN1_ITGA9 | THBS1_PTPRJ | C3_C5AR2 | FN1_ITGA4 | CD38_PECAM1 | APOD_LEPR | CD2_CD48 |
| COL18A1_GPC4 | COL1A1_ITGA11 | SEMA3E_PLXND1 | ADAM15_ITGB1 | PLTP_ABCA1 | SERPING1_LRP1 | CCL2_ACKR1 | EDN1_EDNRA | NID1_ITGB3 | IFNL1_IL10RB |
| NCAM1_GFRA1 | SFRP1_FZD6 | MMP7_CD151 | TGFB2_ENG | TNFSF12_CD163 | COL4A1_ITGAV | TGFB1_ACVRL1 | FN1_ITGB3 | FN1_COL13A1 | GNAI2_MTNR1B |
| COL3A1_DDR2 | INHBA_ENG | A2M_LRP1 | CALR_ITGAV | PDGFB_PDGFRB | EDIL3_ITGB3 | LAMB1_ITGA6 | VEGFB_FLT1 | SLPI_PLSCR4 | APOD_LEPR |
| COL4A1_ITGA1 | BMP7_ENG | TGM2_ITGB1 | CCL2_CCR10 | COL4A2_CD93 | CP_SLC40A1 | BMP2_ENG | FN1_SDC2 | FN1_ITGA5 | NID1_ITGB3 |
| WNT3_FZD7 | MDK_ITGA4 | WNT9B_FZD4 | COL4A1_ITGAV | ICAM3_CD209 | COL4A2_CD93 | GLG1_SELE | NAMPT_INSR | VCAN_EGFR | SERPING1_SELE |
| C1QB_LRP1 | CD177_PECAM1 | THBS1_ITGA6 | TSLP_IL7R | CCN1_ITGB2 | FN1_ITGB1 | NTF3_NTRK2 | C3_ITGAX | CCN1_SDC4 | FN1_ITGA5 |
| ADAM9_ITGB5 | SERPINE1_LRP2 | THBS2_NOTCH3 | PGF_FLT1 | CD209_ICAM3 | CX3CL1_ITGB3 | IGFBP4_LRP6 | A2M_LRP1 | CCN2_LRP6 | NPB_NPBWR2 |
| TGFB3_ENG | NCR3LG1_NCR3 | COL18A1_ITGB1 | C3_CD46 | FN1_ITGB1 | FN1_ITGB3 | MRC1_PTPRC | ANXA1_FPR1 | THBS2_ITGA4 | CCN2_LRP6 |
| TGFB2_ENG | FNDC5_ITGAV | COL1A1_ITGB1 | DCN_TLR4 | MDK_SDC4 | C5_C5AR1 | PTGS2_CAV1 | THBS1_ITGA2B | BMP2_BMPR1A | CD52_SIGLEC10 |
| EDN1_KEL | FGF18_FGFR2 | BGN_FGFR3 | TGFB1_TGFBR3 | THBS1_ITGA4 | SEMA6A_PLXNA4 | TNFSF12_CD163 | MFGE8_ITGAV | COL1A2_ITGA11 | LPL_GPIHBP1 |
| MFGE8_ITGB3 | BGN_TLR2 | COL1A2_CD93 | SERPING1_SELE | IL1B_IL1R1 | EFNA3_EPHA4 | CXCL12_ACKR3 | THBS1_ITGA6 | IGFBP4_FZD8 | JAM3_JAM2 |
| PGF_FLT1 | AMH_EGFR | NODAL_ACVR1C | PECAM1_CD38 | LTBP1_ITGB5 | FN1_SDC2 | PDGFB_PDGFRB | COL1A2_CD93 | MICB_KLRK1 | COL1A2_ITGA11 |
| MFGE8_PDGFRB | COL1A1_DDR2 | COL1A1_GP6 | CSF1_CSF1R | COL1A2_ITGB1 | SEMA3E_PLXND1 | THBS1_ITGA4 | SERPINF1_PLXDC1 | CCL2_ACKR1 | IGFBP4_FZD8 |
| C3_LRP1 | CXCL12_CXCR4 | BMP10_ENG | SELE_CD44 | VEGFB_FLT1 | TIMP1_CD63 | FN1_ITGB3 | BSG_SLC16A1 | BMP2_ENG | DLL3_NOTCH3 |
| CXCL12_ITGB1 | CCL4_CCR1 | ADAM17_NOTCH1 | TGFB2_TGFBR2 | SEMA6D_PLXNA1 | HLA-A_APLP2 | EDN1_EDNRA | NGF_NGFR | IGFBP4_LRP6 | CFH_ITGAM |
| NID1_ITGB3 | LAMB1_ITGB1 | THBS1_LRP1 | BMP2_BMPR1A | LAMA1_ITGA1 | SPON1_APP | COL1A2_ITGB3 | ADAM17_NOTCH1 | ANGPTL2_ITGB1 | FN1_ITGA4 |
| SERPING1_SELE | ADAM9_ITGB1 | FN1_TNFRSF11B | COL1A2_ITGA11 | INHBA_TGFBR3 | CADM3_NECTIN1 | FN1_SDC2 | EFNB2_RHBDL2 | PTGS2_CAV1 | CCL2_ACKR1 |
| FN1_COL13A1 | FN1_ITGA2 | ICAM3_ITGAL | CCL2_ACKR1 | CXCL8_SDC1 | BSG_SLC16A1 | A2M_LRP1 | FN1_PLAUR | TNFSF12_CD163 | SPON2_ITGB1 |
| FN1_ITGA2B | ADAM15_ITGA5 | DLL1_NOTCH3 | BMP2_ENG | THBS1_SDC1 | GNAS_ADRB2 | NAMPT_INSR | THBS1_ITGB3 | MRC1_PTPRC | BMP2_ENG |
| WNT2_FZD5 | COL4A5_CD93 | COL18A1_KDR | PLAU_PLAUR | SEMA6A_PLXNA4 | CALR_LRP1 | ANXA1_FPR1 | RGMA_NEO1 | ANXA1_DYSF | IGFBP4_LRP6 |
| TNC_ITGB3 | ANXA1_FPR3 | FN1_PLAUR | TFPI_VLDLR | CCL19_CCR7 | FN1_CD44 | FN1_MAG | GNAI2_EGFR | THBS1_ITGA4 | CCL5_CCR4 |
| CEACAM1_SELE | DLL4_NOTCH3 | THBS1_ITGB3 | ANGPTL2_ITGB1 | ICAM4_ITGB1 | NECTIN1_CADM3 | THBS1_ITGA6 | CXCL14_CXCR4 | FN1_ITGB3 | MRC1_PTPRC |
| ZP3_MERTK | MUC7_SELL | PDGFD_PDGFRB | PTGS2_CAV1 | CCN2_ITGAM | NID1_ITGB3 | COL1A2_ITGA2 | MMRN2_CD93 | COL1A2_ITGB3 | CXCL12_ACKR3 |

| | | | | | | | | | |
|---|---|---|---|---|---|---|---|---|---|
| FGG_ITGB2 | FN1_NT5E | COL8A1_ITGA1 | TNFSF12_CD163 | A2M_LRP1 | FN1_COL13A1 | BGN_FGFR3 | CCN2_ITGA5 | FN1_SDC2 | THBS1_ITGA4 |
| TGFB2_TGFBR2 | BGN_TLR4 | NID1_PTPRF | LGALS3BP_ITGB1 | COL1A1_DDR2 | FGF1_NRP1 | COL1A2_CD93 | CCN1_ITGB5 | SEMA3E_PLXND1 | FN1_ITGB3 |
| CD14_ITGB1 | HSPG2_ITGB1 | COL18A1_GPC4 | CD38_PECAM1 | NAMPT_INSR | WNT7B_LRP5 | BSG_SLC16A1 | COL1A1_ITGA11 | A2M_LRP1 | VEGFB_FLT1 |
| CCN1_ITGB3 | SEMA6D_PLXNA1 | JAG1_NOTCH1 | C1QA_CSPG4 | DCN_TLR4 | FN1_ITGA5 | THBS1_SDC4 | INHBA_ENG | TGM2_ITGB1 | NECTIN2_CD96 |
| IGFBP4_FZD8 | CXCL12_ITGA5 | COL1A1_CD36 | EDN1_EDNRA | TGM2_SDC4 | AZGP1_ITGAV | FN1_ITGA8 | CP_SLC40A1 | THBS1_ITGA6 | COL1A2_ITGB3 |
| GAS6_MERTK | FCN2_LRP1 | THBS1_ITGB1 | VEGFB_FLT1 | THBS1_TNFRSF11B | GPI_AMFR | THBS1_LRP1 | COL1A1_DDR2 | THBS2_NOTCH3 | FN1_SDC2 |
| BMP2_ENG | CCN1_ITGA5 | NECTIN2_NECTIN2 | PROS1_AXL | KITLG_KIT | HLA-F_LILRB2 | FN1_TNFRSF11B | NID1_COL13A1 | COL18A1_ITGB1 | A2M_LRP1 |
| IGFBP4_LRP6 | FN1_ITGAV | SEMA3F_NRP1 | ANGPT1_ITGB1 | BMP6_ACVR1 | CCN1_ITGB3 | TNFSF10_TNFRSF10A | BMP6_ACVR1 | COL1A1_ITGB1 | NAMPT_INSR |
| MRC1_PTPRC | CCN2_IGF2R | CCN2_ITGA5 | ANXA1_FPR1 | PDGFB_PDGFRA | FN1_ITGB8 | DKK2_KREMEN1 | FBN1_ITGA5 | COL1A2_CD93 | ANXA1_FPR1 |
| LGALS3BP_ITGB1 | MMP9_LRP1 | COL1A1_ITGA11 | PVR_TIGIT | THBS1_ITGA6 | APP_LRP1 | WNT5A_MCAM | FN1_ITGB6 | BSG_SLC16A1 | TGM2_ITGB1 |
| CXCL12_ACKR3 | COL4A2_CD93 | LAMA5_BCAM | MFNG_NOTCH1 | COL1A2_ITGA2 | APLN_APLNR | TNC_ITGA7 | AFDN_F11R | CCL5_CCR1 | THBS1_ITGA6 |
| PDGFB_PDGFRB | FN1_ITGB1 | THBS1_PTPRJ | TGM2_ITGB1 | COL1A1_ITGB1 | COL4A1_ITGB1 | FN1_PLAUR | TNC_EGFR | FBN1_ITGAV | THBS2_NOTCH3 |
| EDN1_EDNRA | LGI2_ADAM22 | CCN2_LRP1 | COL18A1_ITGB1 | CXCL12_ITGA5 | APOE_SCARB1 | THBS1_ITGB3 | APLN_APLNR | CD48_CD2 | COL1A1_ITGB1 |
| COL1A2_ITGB3 | COL1A2_ITGB1 | INHBA_ENG | COL1A1_ITGB1 | THBS1_ITGA3 | NID1_PTPRF | VWF_TNFRSF11B | ADAM15_ITGA5 | CCL5_ACKR1 | COL18A1_ITGB1 |
| VTN_ITGB3 | HRAS_INSR | FGB_ITGB1 | BSG_SLC16A1 | RELN_ITGA3 | COL1A1_CD44 | ADGRE5_CD55 | DLL4_NOTCH3 | THBS1_LRP1 | BGN_FGFR3 |
| A2M_LRP1 | EFNB2_EPHA4 | BMP7_ENG | NECTIN2_NECTIN3 | VTN_TNFRSF11B | NECTIN1_NECTIN1 | CD34_SELE | ANXA1_FPR3 | THBS2_ITGB1 | COL1A2_CD93 |
| THBS1_ITGA2B | COL18A1_ITGA5 | LPL_VLDLR | HP_CD163 | GNAI2_CAV1 | | CCN2_ITGA5 | CXCL12_ITGA5 | CEL_CXCR4 | SERPINF1_PLXDC1 |
| ANGPT1_ITGA5 | FGF4_FGFR3 | WNT3A_FZD5 | TIGIT_PVR | COL1A1_ITGA5 | | VCAM1_ITGB2 | FGF1_CSPG4 | COL18A1_KDR | BSG_SLC16A1 |
| COL18A1_ITGB1 | TGFB1_ENG | CD274_PDCD1 | SELPLG_SELE | INHBA_ACVR1 | | LAMA5_BCAM | DCN_TLR2 | FN1_PLAUR | HP_CD163 |
| GNAI2_CCR5 | TIMP1_CD63 | CCN2_NTRK1 | TNC_ITGAV | FN1_ITGA5 | | THBS1_PTPRJ | CCN1_ITGA5 | C1QA_CD33 | SELPLG_SELE |
| BSG_SLC16A1 | COL4A1_ITGB8 | BGN_TLR2 | SEMA3F_NRP2 | FGF20_FGFR1 | | PLTP_ABCA1 | FN1_ITGAV | CXCL16_CXCR6 | CD48_CD2 |
| SELPLG_SELE | MDK_LRP1 | THBS1_TNFRSF11B | CALR_ITGA3 | FN1_ITGA2 | | HBEGF_CD9 | CCN2_IGF2R | COL18A1_GPC4 | FN1_ITGA8 |
| TNC_ITGAV | MMRN2_CLEC14A | THBS1_ITGA3 | HBEGF_ERBB2 | CCN2_LRP6 | | INHBA_ENG | COL4A2_CD93 | CXCL12_ITGAV | THBS1_LRP1 |
| HBEGF_ERBB2 | EFNB2_PECAM1 | TGM2_ITGB3 | COL18A1_KDR | CCN2_ITGB2 | | BMP7_ENG | TNFSF14_LTBR | JAG1_NOTCH1 | FN1_TNFRSF11B |
| COL18A1_KDR | COL1A1_ITGA5 | PSEN1_NOTCH1 | FN1_PLAUR | FN1_PLAUR | | CD55_ADGRE5 | FN1_ITGB1 | THBS1_ITGB1 | WNT5A_MCAM |
| THBS1_ITGB3 | FN1_ITGB8 | FN1_ITGB6 | C1QA_CD33 | THBS1_ITGB3 | | FBN1_ITGB3 | COL18A1_ITGA5 | COL1A1_CD36 | LGALS1_CD69 |
| COL18A1_GPC4 | SPON2_ITGA5 | LAMB1_ITGB1 | CCL2_CCR1 | FN1_NT5E | | FGF18_FGFR2 | TIMP1_CD63 | MMRN2_CD93 | COL18A1_KDR |
| ADAM17_ITGA5 | AFDN_EPHB6 | GDF2_ACVRL1 | SEMA3C_NRP2 | COL4A1_ITGB1 | | THBS1_SDC1 | EFEMP1_EGFR | THBS2_ITGA6 | FN1_PLAUR |
| FGF17_FGFR3 | COL4A1_ITGB1 | SEMA3D_NRP1 | THBS1_ITGB1 | COL8A1_ITGA1 | | IL32_ITGB3 | BMP6_BMPR1A | CCN2_ITGA5 | THBS1_ITGB3 |
| IL33_IL1RL1 | MCAM_MCAM | CCN2_ITGB2 | MUC1_SIGLEC9 | CCN1_ITGB3 | | GNAS_PTGDR | HRAS_CAV1 | COL1A1_ITGA11 | LRFN5_LRFN5 |
| F13A1_ITGA4 | COL3A1_DDR2 | DLL4_NOTCH3 | SERPINF1_PLXDC2 | COL3A1_DDR2 | | PTPRC_MRC1 | EFNB2_EPHB6 | IL16_CD4 | CXCL12_ITGAV |
| CCN2_ITGA5 | COL4A1_ITGA1 | FN1_NT5E | IL33_IL1RL1 | VTN_PLAUR | | COL1A1_DDR2 | COL4A1_CD93 | CCN2_LRP1 | JAG1_NOTCH1 |
| LAMA5_BCAM | FN1_ITGA9 | BGN_TLR4 | MMRN2_CD93 | VWF_TNFRSF11B | | FGF1_FGFR2 | MMRN2_CLEC14A | PLTP_ABCA1 | THBS1_ITGB1 |

| | | | | | | | | | |
|---|---|---|---|---|---|---|---|---|---|
| GRN_TNFRSF1B | C3_C3AR1 | HSPG2_ITGB1 | COL1A1_ITGA11 | COL4A1_ITGA1 | | CXCL8_ACKR1 | EFNB2_PECAM1 | GRN_TNFRSF1B | F13A1_ITGA4 |
| INHBA_ENG | FSTL1_DIP2A | HSPG2_COL13A1 | NPW_NPBWR2 | | | THBS1_ITGA3 | COL1A1_ITGA5 | CXCL12_ITGA4 | CD58_CD2 |
| CP_SLC40A1 | PSEN1_NOTCH3 | FN1_ITGAV | GRN_TNFRSF1B | | | TGM2_ITGB3 | ICAM1_ITGAX | INHBA_ENG | CD34_SELP |
| BMP7_ENG | COL4A1_ITGAV | CCN2_IGF2R | BMP7_ENG | | | DCN_EGFR | JAG1_NOTCH3 | BMP7_ENG | CCN2_ITGA5 |
| C4B_CR1 | PLA2G10_PLA2R1 | JAG1_CD46 | TNFSF13_TNFRSF1A | | | FBN1_ITGA5 | PTPRK_PTPRK | FBN1_ITGB3 | THBS1_PTPRJ |
| FBN1_ITGB3 | PGF_FLT1 | MMP9_LRP1 | IL32_ITGB3 | | | LGALS1_PTPRC | CX3CL1_ITGAV | GNAI2_TBXA2R | CCN2_LRP1 |
| THBS1_SDC1 | MFGE8_PDGFRB | MYL9_CD69 | BGN_TLR2 | | | CTHRC1_FZD6 | COL4A1_ITGB1 | THBS1_SDC1 | GRN_TNFRSF1B |
| IL32_ITGB3 | NPS_NPSR1 | CCN2_FGFR2 | TNF_TNFRSF1B | | | FN1_ITGA2 | COL4A1_ITGA1 | PTPRC_MRC1 | INHBA_ENG |
| BGN_TLR2 | LTBP1_ITGB5 | FN1_ITGB1 | SLAMF6_SLAMF6 | | | ADAM15_ITGA5 | | CCL5_CCR5 | CXCL12_ITGA4 |
| SERPINC1_LRP1 | JAM2_JAM3 | COL1A2_ITGB1 | SEMA3G_NRP2 | | | THBS1_CD47 | | CXCL8_ACKR1 | PLTP_ABCA1 |
| PTPRC_MRC1 | ICAM5_ITGB2 | HRAS_INSR | SEMA6D_TYROBP | | | LGALS3_LAG3 | | CXCL12_CXCR4 | BMP7_ENG |
| SLIT2_SDC1 | EDN2_EDNRA | SEMA7A_ITGA1 | TGM2_ITGB3 | | | FN1_NT5E | | DCN_EGFR | CD177_PECAM1 |
| TGM2_ITGB3 | FN1_CD44 | LAMA1_ITGA1 | DCN_EGFR | | | BGN_TLR4 | | COL4A1_ITGA1 | LPL_VLDLR |
| CLCF1_CRLF1 | PECAM1_CD38 | COL18A1_ITGA5 | TGFB1_TGFBR2 | | | GDF11_ACVR2B | | FBN1_ITGA5 | FBN1_ITGB3 |
| TGFB1_TGFBR2 | FN1_ITGA2B | CCN2_ITGAM | GNAI2_ADORA1 | | | CD14_ITGA4 | | TGFB1_TGFBR2 | IL32_ITGB3 |
| CCL4_CCR1 | CCN3_NOTCH1 | TIMP1_CD63 | ARF1_INSR | | | PDGFC_PDGFRB | | ADAM9_ITGB1 | BGN_TLR2 |
| ADAM9_ITGB1 | FN1_ITGA5 | COL4A1_ITGB8 | LAMB1_ITGB1 | | | ICAM2_ITGB2 | | CCN2_ITGB2 | PTPRC_MRC1 |
| ADAM15_ITGA5 | TGFB2_TGFBR2 | SLIT1_SDC1 | ADAM9_ITGB1 | | | CXCL8_SDC2 | | DLL4_NOTCH3 | WNT8A_FZD4 |
| DLL4_NOTCH3 | FN1_ITGA4 | COL4A1_CD93 | BGN_TLR4 | | | DCN_TLR2 | | CXCL12_CD4 | CXCL8_ACKR1 |
| ANXA1_FPR3 | GRN_SORT1 | COL1A1_ITGA5 | LY9_LY9 | | | FN1_ITGAV | | DLK1_NOTCH3 | CXCL12_CXCR3 |
| CCL8_CCR1 | LAMB1_ITGA6 | HSPG2_ITGA2 | ICAM2_ITGB2 | | | DCN_MET | | CXCL12_ITGA5 | THBS1_ITGA3 |
| FN1_NT5E | PTGS2_CAV1 | JAG1_NOTCH3 | DCN_TLR2 | | | C3_C5AR2 | | EFNB3_EPHA4 | CXCL12_CXCR4 |
| CXCL12_ITGA5 | NGF_SORT1 | FN1_ITGB8 | DCN_MET | | | EDIL3_ITGB3 | | CXCL8_SDC2 | DCN_EGFR |
| PDGFC_PDGFRB | VWF_ITGA2B | PTPRK_PTPRK | C3_C5AR2 | | | COL4A2_CD93 | | DCN_TLR2 | TGM2_ITGB3 |
| L1CAM_ITGA5 | CD38_PECAM1 | COL4A1_ITGB1 | IL13_IL13RA1 | | | EFNB2_EPHA4 | | FN1_ITGAV | FBN1_ITGA5 |
| ADAM23_ITGA5 | PDGFB_PDGFRB | CXCL12_ITGB3 | ANXA2_ROBO4 | | | TGFB1_ENG | | IL1RN_IL1R1 | LAMB1_ITGB1 |
| DCN_TLR2 | FN1_ITGB3 | MCAM_MCAM | ADAM28_ITGA4 | | | ADAM15_ITGB3 | | CCN2_IGF2R | RAET1E_KLRK1 |
| CCN1_ITGA5 | FN1_SDC2 | COL3A1_DDR2 | IGSF11_VSIR | | | TIMP1_CD63 | | JAG1_CD46 | KNG1_PLAUR |
| FN1_ITGAV | A2M_LRP1 | COL4A1_ITGA1 | FN1_ITGB1 | | | COL4A1_ITGB8 | | AMBN_CD63 | CCN2_ITGB2 |
| JAG1_CD46 | NAMPT_INSR | JAG1_NOTCH1 | PLAU_ST14 | | | CCN2_ITGAM | | MMP9_LRP1 | ADAM15_ITGA5 |
| COL4A2_CD93 | ANGPT1_ITGA5 | NECTIN2_NECTIN2 | F13A1_ITGB1 | | | ICAM3_ITGB2 | | CCN2_FGFR2 | CXCL12_CD4 |
| FN1_ITGB1 | MFGE8_ITGAV | GRN_SORT1 | COL1A2_ITGB1 | | | CCL2_CCR5 | | FN1_ITGB1 | THBS1_CD47 |
| COL1A2_ITGB1 | TGM2_ITGB1 | CCN1_ITGA5 | EFNB2_EPHA4 | | | COL4A1_CD93 | | COL1A2_ITGB1 | BGN_TLR4 |
| LAMA1_ITGA1 | THBS2_NOTCH3 | PODXL2_SELL | COL18A1_ITGA5 | | | DKK2_LRP5 | | CCL19_CCR7 | FN1_NT5E |
| COL18A1_ITGA5 | COL1A1_ITGB1 | CCN2_IGF2R | TGFB1_ENG | | | EFNB2_PECAM1 | | COL18A1_ITGA5 | HSPG2_ITGB1 |

| | | | | | | | | | |
|---|---|---|---|---|---|---|---|---|---|
| TGFB1_ENG | BGN_FGFR3 | SEMA3B_NRP1 | FBN1_ITGB1 | | | C1QB_CD33 | | TGFB1_ENG | CXCL12_ITGA5 |
| ADAM15_ITGB3 | COL18A1_ITGB1 | CCN2_ITGA5 | C1QA_CD93 | | | BSG_SELE | | ADAM15_ITGB3 | MMP2_SDC2 |
| TIMP1_CD63 | COL1A2_CD93 | CCN1_ITGB5 | COL4A1_CD93 | | | HSPG2_PTPRS | | TIMP1_CD63 | PDGFC_PDGFRB |
| ADAM17_ITGB1 | SERPINF1_PLXDC1 | PDGFA_PDGFRA | MMRN2_CLEC14A | | | RSPO3_LRP6 | | TGFB3_TGFBR2 | DCN_TLR2 |
| BMP6_BMPR1A | TGFB1_ITGAV | COL4A1_ITGAV | PTGS2_PTGDR2 | | | FGF1_NRP1 | | FBN1_ITGB1 | CCN1_ITGA5 |
| NXPH3_NRXN2 | FN1_TNFRSF11B | GAS6_AXL | SERPING1_SELP | | | COL1A1_ITGA5 | | COL4A1_CD93 | FN1_ITGAV |
| C1QA_CD93 | NTS_SORT1 | CCN2_LRP1 | BSG_SELE | | | NTN1_MCAM | | EFNB2_PECAM1 | CCN2_IGF2R |
| COL4A1_CD93 | FN1_PLAUR | CD34_SELL | COL4A1_ITGB1 | | | GNAS_VIPR1 | | C1QB_CD33 | DCN_MET |
| MMRN2_CLEC14A | PDGFD_PDGFRB | CD38_PECAM1 | TNFSF18_TNFRSF18 | | | GNAI2_C5AR1 | | VCAN_ITGB1 | JAG1_CD46 |
| EFNB2_PECAM1 | NID1_PTPRF | JAG1_NOTCH2 | ANXA2_TLR2 | | | DKK2_LRP6 | | COL1A1_ITGA5 | MYL9_CD69 |
| BSG_SELE | VWF_TNFRSF11B | DLL4_NOTCH1 | COL3A1_DDR2 | | | HSPG2_ITGA2 | | RAET1L_KLRK1 | COL4A2_CD93 |
| COL1A1_ITGA5 | COL1A1_CD36 | LTBP1_ITGB5 | COL4A1_ITGA1 | | | CALR_SCARF1 | | HSPG2_ITGA2 | FN1_ITGB1 |
| IL16_KCNJ15 | ARF1_PLD2 | FGF5_FGFR1 | | | | EFNA3_EPHA3 | | F13A1_ITGA9 | PLAU_ST14 |
| F13A1_ITGA9 | NECTIN2_NECTIN2 | CD226_NECTIN2 | | | | JAG1_NOTCH3 | | CALR_SCARF1 | F13A1_ITGB1 |
| JAG1_NOTCH3 | COL4A6_CD93 | PDGFB_PDGFRA | | | | FN1_ITGB8 | | JAG1_NOTCH3 | COL1A2_ITGB1 |
| FN1_ITGB8 | ADAM2_ITGA9 | THBS2_NOTCH3 | | | | TNFSF10_TNFRSF10B | | MIF_CXCR4 | CCL19_CCR7 |
| SPON2_ITGA5 | SEMA3F_NRP1 | LAMC2_ITGB1 | | | | EFNB2_EPHB4 | | CXCL12_ITGB3 | COL18A1_ITGA5 |
| COL4A1_ITGB1 | IL16_CD4 | EFNB2_RHBDL2 | | | | CXCL12_ITGB3 | | GZMB_IGF2R | TGFB1_ENG |
| CXCL12_ITGB3 | INHBA_ENG | BMP2_BMPR1B | | | | MCAM_MCAM | | CCL8_ACKR2 | CCN2_ITGAM |
| COL3A1_DDR2 | CP_SLC40A1 | GDF6_BMPR2 | | | | COL1A1_CD44 | | CXCL1_ACKR1 | TIMP1_CD63 |
| COL1A2_GP6 | BMP7_ENG | CTHRC1_FZD6 | | | | GZMB_IGF2R | | | FBN1_ITGB1 |
| COL4A1_ITGA1 | NCR3LG1_NCR3 | SEMA3D_NRP1 | | | | | | | EFNB2_EPHB6 |
| | AGRN_DAG1 | CCN1_SDC4 | | | | | | | COL4A1_CD93 |
| | BGN_TLR2 | JAG1_NOTCH3 | | | | | | | EFNB2_PECAM1 |
| | COL1A1_DDR2 | NECTIN2_CD226 | | | | | | | GAL_GALR2 |
| | FN1_ITGB6 | CCN2_LRP6 | | | | | | | BSG_SELE |
| | ARF1_INSR | CCN2_ITGB2 | | | | | | | COL1A1_ITGA5 |
| | ADAM9_ITGB1 | DLL4_NOTCH3 | | | | | | | JAG1_NOTCH3 |
| | CD80_CD274 | EFNB2_EPHB4 | | | | | | | FN1_ITGB8 |
| | FN1_ITGA2 | COL4A1_ITGB1 | | | | | | | MIF_CXCR4 |
| | APLN_APLNR | CALCB_CALCR | | | | | | | COL4A1_ITGB1 |
| | FGF20_FGFR4 | LRPAP1_SORL1 | | | | | | | ANXA2_TLR2 |
| | DLL4_NOTCH3 | PODXL_SELL | | | | | | | CXCL12_ITGB3 |
| | FN1_NT5E | | | | | | | | MCAM_MCAM |
| | BGN_TLR4 | | | | | | | | COL3A1_DDR2 |

| | | | | | | | | | |
|---|---|---|---|---|---|---|---|---|---|
| | HSPG2_ITGB1 | | | | | | | | COL4A1_ITGA1 |
| | SEMA3F_PLXNA3 | | | | | | | | |
| | CLEC4G_LAG3 | | | | | | | | |
| | FN1_ITGAV | | | | | | | | |
| | CCN2_IGF2R | | | | | | | | |
| | MMP9_LRP1 | | | | | | | | |
| | CD274_CD80 | | | | | | | | |
| | COL4A2_CD93 | | | | | | | | |
| | RELN_ITGB1 | | | | | | | | |
| | PGF_NRP1 | | | | | | | | |
| | LAMA2_ITGB1 | | | | | | | | |
| | FN1_ITGB1 | | | | | | | | |
| | COL1A2_ITGB1 | | | | | | | | |
| | HRAS_INSR | | | | | | | | |
| | LAMA1_ITGA1 | | | | | | | | |
| | COL18A1_ITGA5 | | | | | | | | |
| | FGG_ITGB1 | | | | | | | | |
| | TGFB1_ENG | | | | | | | | |
| | TIMP1_CD63 | | | | | | | | |
| | COL4A1_ITGB8 | | | | | | | | |
| | TNFSF4_TNFRSF4 | | | | | | | | |
| | LGALS1_ITGB1 | | | | | | | | |
| | LAMA2_ITGA7 | | | | | | | | |
| | MMRN2_CD248 | | | | | | | | |
| | COL4A1_CD93 | | | | | | | | |
| | MMRN2_CLEC14A | | | | | | | | |
| | EFNB2_PECAM1 | | | | | | | | |
| | COL1A1_ITGA5 | | | | | | | | |
| | ANGPT1_TIE1 | | | | | | | | |
| | AZGP1_ITGAV | | | | | | | | |
| | AGRN_LRP4 | | | | | | | | |
| | HSPG2_ITGA2 | | | | | | | | |
| | JAG1_NOTCH3 | | | | | | | | |
| | FN1_ITGB8 | | | | | | | | |
| | SPON2_ITGA5 | | | | | | | | |
| | COL4A1_ITGB1 | | | | | | | | |

| | |
|---|---|
| | SLIT3_ROBO2 |
| | COL3A1_DDR2 |
| | NID1_ITGAV |
| | COL4A1_ITGA1 |

**Supplementary Table S2: Ligand-Receptor Pairs, Neuronal-to-Tumour Communities**

| Midbrain | PonsA | PonsB | PonsC | PonsD | MedullaA | MedullaB | MedullaC | SpinalCordA | SpinalCordB |
|---|---|---|---|---|---|---|---|---|---|
| SST_SSTR5 | FGF19_FGFR1 | SLITRK2_PTPRD | THBS1_CD36 | LGI3_ADAM23 | EFNA1_EPHA5 | LGI3_ADAM23 | LRPAP1_LDLR | NLGN1_NRXN1 | CCN1_ITGAM |
| PTPRM_PTPRM | LGI3_ADAM23 | ADCYAP1_ADCYAP1R1 | WNT5A_MCAM | ADCYAP1_ADCYAP1R1 | NLGN2_NRXN3 | ITGB2_THY1 | ADCYAP1_ADCYAP1R1 | BMP2_BMPR2 | VEGFC_NRP2 |
| DLK1_NOTCH2 | FGF19_FGFR3 | FGF1_FGFR3 | ADCYAP1_SCTR | IFNA16_IFNAR2 | TIMP3_KDR | NLGN3_NRXN1 | HRAS_CAV1 | ADCYAP1_ADCYAP1R1 | CCN1_ITGAV |
| L1CAM_EPHB2 | L1CAM_ERBB3 | SHH_GPC5 | LGI3_ADAM22 | PGF_FLT1 | CCN2_LRP1 | MDK_ALK | WNT5A_MCAM | SLIT1_ROBO2 | EDN1_EDNRA |
| NLGN4X_NRXN1 | LGI1_ADAM23 | DLK1_NOTCH2 | PTPRM_PTPRM | LGI3_ADAM22 | MDK_LRP1 | ADGRB1_RTN4RL2 | GZMB_CHRM3 | EPCAM_EPCAM | CCN1_ITGA5 |
| PTHLH_PTH1R | PTPRM_PTPRM | L1CAM_ERBB2 | FGF2_FGFR1 | NTF3_NTRK1 | FN1_ITGA9 | PTN_ALK | LGI1_ADAM23 | SST_SSTR2 | ADCYAP1_VIPR1 |
| OMG_LINGO1 | CD52_SIGLEC10 | WNT5A_LRP5 | CADM1_CRTAM | LGI1_ADAM23 | EFNA1_EPHA5 | | SLIT3_ROBO2 | NRG1_ERBB4 | |
| ADCYAP1_VIPR2 | PLA2G2A_ITGB3 | CALCB_CALCRL | DLL4_NOTCH3 | LGI2_ADAM23 | AGRN_ATP1A3 | | WNT7A_FZD1 | PDYN_OPRM1 | |
| LGI4_ADAM11 | L1CAM_ERBB2 | FGF5_FGFR3 | TAC4_TACR3 | LGI3_STX1A | SPP1_ITGB3 | | FGF17_FGFR3 | PENK_OGFR | |
| L1CAM_ALCAM | FGF19_FGFR2 | GNAI2_ADRA2B | CXCL5_ACKR1 | ADAM23_ITGA4 | SLIT3_ROBO1 | | LGI2_ADAM23 | NLGN4X_NRXN1 | |
| LGI1_ADAM11 | LGI1_ADAM11 | GAL_GALR3 | FGF17_FGFR1 | LGI1_ADAM11 | PSEN1_NOTCH4 | | WNT7B_FZD8 | RPH3A_NRXN1 | |
| LGI1_ADAM22 | L1CAM_EGFR | SFRP1_FZD2 | CRTAM_CADM1 | LGI1_RTN4R | AFDN_EPHA7 | | NPY_PRLHR | CCK_CCKBR | |
| INHBA_BAMBI | BDNF_NGFR | ADCYAP1_VIPR2 | INHBA_ACVR1B | ADCYAP1_VIPR2 | A2M_LRP1 | | ADCYAP1_VIPR2 | PDYN_OPRK1 | |
| ROBO2_ROBO2 | ADAM15_ITGB3 | FGF2_FGFRL1 | NECTIN1_NECTIN4 | IFNW1_IFNAR2 | C1QA_CD93 | | EFNA5_EPHA7 | LGI1_ADAM23 | |
| LGI2_ADAM23 | CALCB_CALCRL | CALCB_RAMP1 | EFNA5_EPHA5 | BMP6_ACVR1 | TGFB3_TGFBR1 | | NXPH2_NRXN1 | PENK_OPRM1 | |
| ALCAM_L1CAM | MMP7_ERBB4 | FGF2_FGFR3 | MCAM_MCAM | LGI1_ADAM11 | FGF2_CD44 | | SEMA3C_NRP1 | WNT4_FZD6 | |
| CADM3_CADM3 | ARF1_CHRM3 | RTN4_RTN4R | LGI1_ADAM22 | ROBO2_ROBO2 | AFDN_EPHB6 | | | DLK1_NOTCH1 | |
| NPY_NPY5R | LGI1_ADAM23 | FGF2_GPC4 | SERPING1_SELE | ADCYAP1_ADCYAP1R1 | GNAI2_ADCY1 | | | FGF13_SCN8A | |
| EFNA5_EPHA5 | IL4_CD53 | BSG_SLC16A1 | SEMA7A_PLXNC1 | LGI1_ADAM23 | GRN_TNFRSF1A | | | NLGN3_NRXN1 | |
| L1CAM_ALCAM | GHRH_VIPR1 | WNT5A_FZD6 | BGN_TLR4 | IL33_IL1RL1 | | | | ARF1_INSR | |
| FN1_ITGB3 | SEMA3C_NRP2 | WNT7B_LRP5 | TGFB1_SDC2 | BMP5_HJV | | | | ROBO2_ROBO2 | |
| L1CAM_ERBB3 | LGI2_ADAM22 | GDF6_BMPR2 | LGI1_ADAM11 | MCAM_MCAM | | | | NXPH1_NRXN1 | |
| RELN_ITGA3 | LGI1_ADAM22 | PDGFC_PDGFRA | C4BPA_LRP1 | LGI1_ADAM22 | | | | LIN7C_HTR2C | |
| LIN7C_HTR2C | CADM1_CADM1 | ROBO2_ROBO2 | LGI1_ADAM23 | LGALS3_LAG3 | | | | NPY_NPY1R | |

| | | | | | | | | |
|---|---|---|---|---|---|---|---|---|
| GZMB_CHRM3 | CRTAM_CADM1 | WNT5A_MCAM | ADCYAP1_VIPR1 | TAC1_TACR1 | | | | GRP_NMBR |
| CCL5_SDC1 | CCL25_ACKR2 | FGF2_SDC2 | IL31_OSMR | CP_SLC40A1 | | | | LRFN5_LRFN5 |
| AGRP_MC4R | SLIT3_ROBO2 | FGG_ITGAV | TFF2_CXCR4 | LRFN5_LRFN5 | | | | NPY_NPY5R |
| PENK_OPRM1 | WNT2_FZD4 | WNT5A_ROR1 | INHBA_BAMBI | LGI1_RTN4R | | | | GZMB_IGF2R |
| PTPRM_PTPRM | CADM1_CRTAM | SEMA4D_PLXNB2 | TNFSF13_SDC2 | CCN1_ITGB3 | | | | L1CAM_ALCAM |
| CALCB_CALCRL | MMP7_CD44 | ADCYAP1_ADCYAP1R1 | LGI1_ADAM22 | CCN1_ITGAM | | | | PENK_OGFR |
| RELN_VLDLR | | NTN4_UNC5A | CCN1_ITGB5 | NPY_NPY1R | | | | DLK1_NOTCH1 |
| SLIT3_ROBO2 | | CCL2_CCR10 | VCAM1_ITGB2 | NTF4_NTRK1 | | | | FGF4_FGFR3 |
| PLA2G2A_ITGA5 | | EFNA5_EPHA2 | ADCYAP1_SCTR | L1CAM_ITGA5 | | | | DLK1_NOTCH2 |
| EFNA5_EPHB6 | | EFNB1_EPHB6 | FGA_TLR4 | L1CAM_ERBB2 | | | | ALCAM_L1CAM |
| EPCAM_EPCAM | | SEMA4D_PLXNB1 | LPL_LRP2 | ADAM23_ITGA5 | | | | LIN7C_HTR2C |
| SST_SSTR3 | | CCL2_CCR5 | ANXA1_DYSF | | | | | CD80_CD274 |
| TNC_ITGB1 | | LRFN5_LRFN5 | RTN4_RTN4R | | | | | RELN_ITGB1 |
| LAMA3_SDC2 | | MCAM_MCAM | SST_SSTR1 | | | | | EFNB3_EPHA4 |
| TNC_ITGA7 | | ADAM9_ITGA3 | BGN_TLR2 | | | | | CD274_CD80 |
| TNC_ITGAV | | CCN1_ITGB3 | LGI2_ADAM23 | | | | | |
| CRH_CRHR2 | | IL3_CSF2RB | SEMA7A_ITGB1 | | | | | |
| L1CAM_ITGA5 | | ADCYAP1_VIPR2 | LGI3_ADAM23 | | | | | |
| ALCAM_L1CAM | | RTN4_TNFRSF19 | THBS1_ITGB3 | | | | | |
| GZMB_IGF2R | | CNTN1_NOTCH2 | ANXA1_FPR2 | | | | | |
| RELN_LRP8 | | | KNG1_PLAUR | | | | | |
| LGI2_ADAM23 | | | RTN4_CNTNAP1 | | | | | |
| CADM1_CADM1 | | | INHBA_ACVR1 | | | | | |
| CALCB_RAMP1 | | | ITGB2_THY1 | | | | | |
| GHRL_GHSR | | | | | | | | |
| PLA2G2A_ITGB3 | | | | | | | | |

**Supplementary Table S3: Ligand-Receptor Pairs, Myeloid-to-Tumour Communities**

| Midbrain | PonsA | PonsB | PonsC | PonsD | MedullaA | MedullaB | MedullaC | SpinalCordA | SpinalCordB |
|---|---|---|---|---|---|---|---|---|---|
| FN1_ITGB8 | VEGFD_ITGA9 | SEMA6D_TREM2 | VTN_ITGA8 | C3_ITGB2 | NCAM1_PTPRZ1 | ADAM23_ITGA5 | CCL3_CCR1 | PTN_PTPRS | SEMA6B_PLXNA2 |
| EFNB3_EPHB3 | GDF5_ACVR2A | C1QB_CD33 | RAET1G_KLRK1 | C3_C3AR1 | HBEGF_CD82 | FN1_ITGB8 | IFNA2_IFNAR2 | C3_ITGB2 | CD84_CD84 |
| CXCL12_CD4 | MYL9_CD69 | COMP_ITGA5 | EFNB1_EPHA4 | NECTIN3_NECTIN3 | OSM_IL6ST | MDK_LRP1 | ZP3_MERTK | EFNB1_EPHB1 | CLEC11A_ITGB1 |

| | | | | | | | | | |
|---|---|---|---|---|---|---|---|---|---|
| GNAI2_LHCGR | IFNG_IFNGR2 | SFTPD_LAIR1 | ARF1_INSR | COL2A1_ITGB1 | SLIT2_APP | SEMA3C_PLXND1 | OMG_RTN4RL1 | EFNB3_EPHB3 | CADM3_CADM3 |
| ICAM5_ITGB2 | FGB_ITGAM | C3_CD19 | EFNA2_EPHA7 | SEMA3C_NRP2 | HMGB1_SDC1 | IL1B_IL1R2 | TGFB1_ITGB8 | LPL_LRP2 | C3_C3AR1 |
| SEMA3C_NRP2 | CRP_OLR1 | | EPHA4_EFNB1 | ADM_CALCRL | ADM_RAMP2 | SERPINE2_LRP1 | SLITRK1_PTPRS | NCAM1_ROBO1 | CHAD_ITGB1 |
| ADM_CALCRL | CD24_SELP | | FGF2_NRP1 | COL1A2_CD44 | IL34_PTPRZ1 | TFPI_LRP1 | A2M_LRP1 | C3_LRP1 | ANGPTL2_ITGB1 |
| C3_ITGB2 | WNT4_FZD6 | | MUC1_SIGLEC9 | HSP90AA1_FGFR3 | BGN_TLR2 | FN1_NT5E | AFDN_F11R | C3_C3AR1 | TGFB3_TGFBR1 |
| ADAM15_ITGB1 | IFNA14_IFNAR1 | | GRN_SORT1 | LRPAP1_LRP1 | CADM1_CADM1 | AMELY_CD63 | AFDN_NECTIN4 | NCAM1_FGFR1 | CCN1_ITGB2 |
| C3_C5AR2 | EREG_EGFR | | C3_NRP1 | SEMA3B_NRP1 | HBEGF_EGFR | CSF1_CSF1R | LTF_LRP1 | TFPI_LRP1 | ICAM1_ITGAX |
| C3_ITGAM | FGA_ITGB2 | | C3_ITGB2 | SPP1_ITGB5 | NXPH3_NRXN2 | PDGFA_PDGFRA | CCN1_ITGAM | EFNB3_EPHB1 | SERPINF1_PLXDC2 |
| IL16_CD4 | ADIPOQ_ADIPOR2 | | SERPINF1_PLXDC2 | EFNA1_EPHA6 | GRN_TNFRSF1A | EFNA2_EPHA7 | FGF2_SDC4 | NTN1_NEO1 | ITGB2_THY1 |
| NPW_NPBWR2 | BMP2_ACVR2A | | CX3CL1_CX3CR1 | NLGN1_NRXN3 | RTN4_CNTNAP1 | CXCL12_ITGA5 | NECTIN1_NECTIN3 | C3_ITGAX | CALR_ITGA3 |
| SIRPB2_CD47 | VEGFD_FLT4 | | GNAI2_CNR1 | SPP1_ITGB3 | SPP1_ITGB1 | CCN2_LRP1 | SEMA4D_MET | C3_CD46 | WNT3A_RYK |
| FGF20_FGFR2 | FGF2_NRP1 | | B2M_LILRB1 | NLGN3_NRXN3 | TNC_ITGB1 | CLCF1_IL6ST | LTF_LRP11 | EFNB3_EPHB2 | SIRPG_CD47 |
| CCN1_ITGB2 | C3_NRP1 | | LTA_LTBR | HLA-A_APLP2 | SLITRK1_PTPRS | BGN_FGFR3 | INHBA_ACVR1 | SLIT3_ROBO4 | CADM3_CADM3 |
| C3_LRP1 | LTB_TNFRSF1A | | F13A1_ITGA4 | CXCL12_CXCR4 | PTPRZ1_NCAM1 | S100A8_TLR4 | NPPA_NPR1 | C1QB_LRP1 | GNAI2_TBXA2R |
| VEGFA_NRP2 | NTS_SORT1 | | C3_C3AR1 | APOD_LEPR | TNC_EGFR | C3_ITGB2 | CCL23_CCR1 | C3_NRP1 | SEMA3F_NRP2 |
| EFEMP1_EGFR | GNAI2_CCR5 | | C3_ITGAM | SLIT2_ROBO2 | NTN4_DCC | C1QA_CD93 | SPON2_ITGA4 | EFNB3_EPHA4 | WNT8A_FZD4 |
| C3_ITGAX | CD28_CD80 | | CXCL12_ITGA4 | SPP1_ITGAV | HSPG2_ITGB1 | SELPLG_SIGLEC5 | EFEMP1_EGFR | EFNB3_EPHB6 | IFNA8_IFNAR1 |
| ANXA1_FPR1 | VCAM1_ITGA9 | | SDC2_PTPRJ | TF_TFR2 | TNC_CNTN1 | TGFB2_ACVR1 | LGALS3BP_ITGB1 | CADM1_NECTIN3 | |
| ICAM5_ITGB2 | SEMA6A_PLXNA4 | | IL2_CD53 | GPC3_CD81 | SLIT2_GPC1 | CADM3_CADM3 | NECTIN1_NECTIN1 | CCL2_ACKR2 | |
| JAML_CXADR | LRPAP1_SORL1 | | CP_SLC40A1 | NTNG1_LRRC4C | SPP1_CD44 | SERPINA1_LRP1 | SEMA5A_MET | EFNB3_EPHB1 | |
| LGALS9_HAVCR2 | VCAM1_ITGB2 | | GNAI2_FPR1 | RGMA_NEO1 | | EFNB2_EPHB2 | AMH_EGFR | CCL2_CCR1 | |
| IFNA8_IFNAR1 | AFDN_NECTIN3 | | CCN1_ITGB2 | SPP1_CD44 | | SEMA4D_PLXNB2 | | TNFSF13_FAS | |
| WNT3A_RYK | ARF1_CHRM3 | | C1QA_CSPG4 | | | BGN_TLR2 | | CCL7_ACKR1 | |
| LPL_GPIHBP1 | RSPO3_SDC4 | | ADAM28_ITGA4 | | | PLAU_PLAUR | | BMP7_BMPR1B | |
| EFNA5_EPHA3 | CD80_CD28 | | LTBP1_ITGB5 | | | TGFB3_TGFBR2 | | VCAM1_ITGA4 | |
| VCAM1_ITGB1 | CD14_ITGA4 | | MDK_NOTCH2 | | | PLTP_ABCA1 | | NECTIN3_CADM1 | |
| | | | NECTIN2_CD96 | | | CCL18_PITPNM3 | | | |
| | | | C3_ITGAX | | | GNAI2_C5AR1 | | | |
| | | | C3_CD46 | | | | | | |
| | | | DLL3_NOTCH2 | | | | | | |
| | | | RELN_LRP8 | | | | | | |
| | | | PVR_CD226 | | | | | | |
| | | | HBEGF_EGFR | | | | | | |
| | | | RELN_ITGA3 | | | | | | |
| | | | DLL1_NOTCH2 | | | | | | |
| | | | NMB_GRPR | | | | | | |

| | | | |
|---|---|---|---|
| | | | CCL5_CCR1 |
| | | | F10_F3 |
| | | | EFNA2_EPHA7 |
| | | | ANGPT1_TIE1 |
| | | | GNAI2_IGF1R |
| | | | APLN_APLNR |
| | | | TGFB1_TGFBR1 |
| | | | CD14_ITGB1 |
| | | | GNAI2_EGFR |
| | | | LILRB4_LAIR1 |
| | | | CD14_ITGA4 |
